# A tomato MIK2-clade receptor is involved in the perception of a *Fusarium*-derived elicitor

**DOI:** 10.1101/2025.05.22.655486

**Authors:** Julian Maroschek, Yvonne Rösgen, Clemens Rössner, Simon Snoeck, Claus Schwechheimer, Cyril Zipfel, Ralph Hückelhoven

## Abstract

Plants continuously monitor their environment to adapt to threats affecting health and fitness. Cell-surface receptors detect conserved microbe-associated molecular patterns (MAMPs) or endogenous immunogenic signals to activate signaling pathways that induce broad-spectrum disease resistance, known as pattern-triggered immunity (PTI).

In *Arabidopsis thaliana*, the leucine-rich repeat receptor kinase MIK2 emerged as a versatile receptor involved in perceiving endogenous SCOOP peptides as well as potential MAMPs derived from *Fusarium* and related fungi. While SCOOP perception appears restricted to Brassicales, the *Fusarium*-derived elicitor triggers PTI-responses across different plant lineages.

Here, we demonstrate that *Fusarium* elicitor-responsiveness and MIK2-clade proteins are conserved among seed plants. We identified a tomato MIK2-clade receptor that shares *At*MIK2 properties in perceiving the *Fusarium* elicitor, but not SCOOPs. Tomato mutants lacking the receptor show compromised PTI-responses to the fungal elicitor and increased susceptibility to *Fusarium oxysporum*.

By integrating AI-assisted structural prediction with experimental data, we identify conserved and variable residues in *Arabidopsis* and tomato MIK2-clade receptors that may underlie shared and lineage-specific recognition specificities. Together, these findings establish MIK2-clade receptors as immune surveillance modules that have undergone lineage-specific diversification, supporting a model in which endogenous danger perception might have evolved from an ancestral role in microbial pattern recognition.

## Introduction

To respond appropriately to environmental changes and potential threats, plants must sense and transmit signals from their surroundings. Pathogenic microbes and pests pose major challenges to plant fitness and survival. Accordingly, plants continuously monitor their environment using membrane-localized pattern recognition receptors (PRRs). These receptors – either receptor kinases (RKs) or receptor proteins (RPs) – recognize conserved microbial molecules collectively termed microbe-associated molecular patterns (MAMPs) (Boller & Felix, 2009; Basu *et al*., 2018) . In addition, plants detect endogenous danger signals known as damage-associated molecular patterns (DAMPs). Primary DAMPs are passively released upon tissue damage caused by pathogens or pests, whereas secondary endogenous danger signals are actively processed and released in response to damage or other stimuli. Owing to their analogy to metazoan cytokines, these secondary danger molecules are commonly referred to as phytocytokines (Gust *et al*., 2017) .

Ligand-perception by PRRs activates conserved signaling pathways that induce broad-spectrum disease resistance, termed pattern-triggered immunity (PTI) (Dodds *et al*., 2024; Zhang *et al*., 2024) . Upon recognition of proteinaceous ligands, PRRs belonging to the class of leucine-rich repeat receptor kinases (LRR-RKs) recruit co-receptors of the SOMATIC EMBRYOGENESIS RECEPTOR-LIKE KINASE (SERK) family, including BRASSINOSTEROID INSENSITIVE 1-ASSOCIATED RECEPTOR KINASE 1 (BAK1) or BAK1-LIKE 1 (BKK1) to form ligand-dependent receptor complexes (DeFalco & Zipfel, 2021) . Receptor-like cytoplasmic kinases (RLCKs), such as BOTRYTIS-INDUCED KINASE 1 (BIK1) and AVRPPHB SUSCEPTIBLE 1(PBS1)-LIKE 1 (PBL1), constitutively associate with the cytoplasmic domains of these PRR complexes (Hailemariam *et al*., 2024) . Following ligand-perception and receptor-activation, the signal is transduced to the cytoplasm, initiating rapid and transient elevations of cytoplasmic free-calcium concentrations ([Ca^2+^]_cyt_), apoplastic reactive oxygen species (ROS) production, and phosphorylation-dependent activation of mitogen-activated protein kinase (MAPK) cascades (Köster *et al*., 2022; Sun & Zhang, 2022; Wu *et al*., 2023) . These events activate transcriptional regulators, including transcription factors (TFs) and chromatin modifiers, leading to extensive transcriptional reprogramming (Bigeard *et al*., 2015; Li *et al*., 2016) . Supported by posttranscriptional, epigenetic, and posttranslational regulators, this reprogramming culminates in a broad array of defense responses (Li *et al*., 2016; Withers & Dong, 2017; Tao *et al*., 2023), including phytohormone production (e.g., ethylene), biosynthesis and secretion of antimicrobial compounds such as phytoalexins, and reinforcement of cell walls through lignin and callose deposition (Wang *et al*., 2021; Ninkuu *et al*., 2022; Jian *et al*., 2024; Molina *et al*., 2024) .

PTI signaling is tightly regulated by positive and negative feedback mechanisms that amplify weak responses or attenuate excessive signaling. One such mechanism involves membrane-localized receptors that perceive phytocytokines. Phytocytokines are released through enzymatic processing, mediate intercellular signaling, and can function in both growth and development as well as immunity, positioning them as key players in the growth-defense trade-off (Rzemieniewski & Stegmann, 2022) .

In most cases, PRRs and phytocytokine receptors are specialized for the perception of either exogenous or endogenous signals. However, rare exceptions exist. One example is the NEMATODE-INDUCED LRR-RLK 1 (NILR1) receptor from *Solanum tuberosum* (potato), which recognizes both the exogenous nematode-associated molecular pattern (NAMP) ascaroside and the endogenous hormone brassinosteroid (Huang *et al*., 2023; Huang *et al*., 2024) . Another example is MALE DISCOVERER 1-INTERACTING RECEPTOR-LIKE KINASE 2 (MIK2) from *Arabidopsis thaliana*. MIK2 serves as the receptor for the SERINE-RICH ENDOGENOUS PEPTIDE (SCOOP) family, which is specific to the order Brassicales and comprises 50 members in *A. thaliana* alone (Hou *et al*., 2021; Rhodes *et al*., 2021; Yang *et al*., 2023a; Snoeck *et al*., 2024) . Aside from a conserved SxS motif present in most SCOOPs, their amino acid sequences are highly variable (Gully *et al*., 2019; Yang *et al*., 2023a; Snoeck *et al*., 2024) . Recent AI-assisted structural modeling and phylogenomic analysis revealed that this conserved motif mediates SCOOP binding to MIK2 (Snoeck *et al*., 2024) . This was further supported by crystallography and cryo-EM structural studies, which also indicated that *N*-glycosylation of MIK2 facilitates interaction with the co-receptor BAK1 (Jia *et al*., 2024; Wu *et al*., 2024) .

Beyond SCOOP perception, MIK2 is indispensable for PTI responses triggered by an elicitor present in extracts of diverse *Fusarium* and related fungal species. These fungal-derived elicitor fractions were enriched by size-exclusion and ion-exchange chromatography and, in the case of *F. oxysporum,* termed ‘Enriched *Fusarium oxysporum* Elicitor’ (EnFOE). EnFOE triggers hallmarks of PTI, including cytosolic calcium influx, ROS burst, MAP kinase activation, and marker gene expression, depending on BIK1, PBL1, and RBOHD. Although the immunogenic molecule in EnFOE remains unidentified, basic analyses indicate a proteinaceous nature (Coleman *et al*., 2021) . Guided by this observation, bioinformatic analyses identified SCOOP-LIKE (SCOOPL) epitopes within *Fusarium* proteins. Synthetic SCOOPL peptides modeled after plant SCOOPs similarly induce MIK2-dependent PTI responses in *Arabidopsis* (Hou *et al*., 2021; Rhodes *et al*., 2021) . However, most SCOOPL sequences are poorly conserved beyond *Fusarium*, whereas comparable elicitor-activity is also detected in extracts from the related genus *Trichoderma* (Coleman *et al*., 2021) . Moreover, the *Fusarium* elicitor induces PTI-like responses in non-Brassicales species, while SCOOP and SCOOPL perception is restricted to this plant order (Coleman *et al*., 2021; Snoeck *et al*., 2024) .

Here we report the identification of an LRR-RK from tomato that shares a phylogenetic origin with *Arabidopsis thaliana* MIK2 (*At*MIK2). This tomato receptor shows similar functions in response to the *Fusarium* elicitor and in resistance to *Fusarium* infection, but does not mediate perception of plant SCOOPs or synthetic *Fusarium* SCOOPL peptides. By combining AI-assisted structural predictions with experimental data, we identified conserved and variable residues in *Arabidopsis* and tomato MIK2-clade receptors that may underpin both shared and lineage-specific immune responses. These findings suggest that the Brassicales-specific role of MIK2 as a SCOOP receptor could have evolved from a more ancient function as a PRR recognizing a fungal MAMP.

## Material and Methods

### Plant lines and growth conditions

For ROS measurements, *Arabidopsis thaliana* Col-0, *Brassica rapa* subsp. *chinensis* (Pak-Choi), *Glycine max* cv. Williams 82, *Solanum lycopersicum* cv. M82 and *Hordeum vulgare* cv. Golden Promise were planted in soil:vermiculite (ratio 8:1), stratified for 2 days (dark; 4°C) and cultivated under long-day conditions (16 hours light : 8 hours dark photoperiod; 20 – 22°C [16°C for *Hordeum vulgare*]; 55% rel. hum.) for 6 – 8 weeks. Tomato CRISPR mutants and wildtype plants for ROS and ethylene measurements were grown for 8 – 12 weeks under the same conditions. *Nicotiana benthamiana* plants were grown under the same conditions, but without stratification.

CRISPR-Cas9 mediated knock-out of *AtMIK2* and *SlMIK2-A* was performed using apoaequorin-expressing *Arabidopsis thaliana* Col-0^AEQ^ (pMAQ2, Marc Knight, University of Durham, UK) or *Solanum lycopersicum* cv. M82, respectively.

For aequorin luminescence measurements, seeds of *A. thaliana* Col-0^AEQ^ wildtype, *mik2*^AEQ^ CRISPR and *fls2*-26^AEQ^ (Ranf *et al*., 2012) mutants were surface-sterilized, stratified for 2 days (dark; 4°C) and grown under long-day conditions (16 hours light : 8 hours dark photoperiod; 22°C) in liquid medium (½ MS plus vitamins, 2.5 g/L sucrose, 1 mM MES, pH 5.7) for 7 days.

For hydroponic growth (Araponics, Belgium), *S. lycopersicum* cv. M82 seeds were surface-sterilized with chlorine gas, placed in seed-holders with solid medium (MS plus vitamins,1 g/L phytagel) and stratified for 2 days (dark, 4°C). Seed-holders were transferred to hydroponic boxes with standard medium (Tocquin *et al*., 2003), covered, and grown under long-day conditions as described above. Medium was replaced weekly, and lids removed two weeks after germination.

### Molecular cloning

Sequences for transient over-expression in *Nicotiana benthamiana* were cloned into GoldenGate modules using appropriate primers and assembled under CaMV35S promotor and terminator control. Gene cassettes were ligated into a GoldenGate-modified pCB302 binary vector. Cloned sequences and used primers are listed in table S1a-b. Proteins for gain-of-function, co-immunoprecipitation, and confocal imaging were fused to C-terminal cMyc, GFP or mCherry tags.

For the apoplastic expression of LRR ectodomains, non-cytoplasmic domains were predicted (PHOBIUS; (Käll *et al*., 2004) ; https://phobius.sbc.su.se/index.html) and amplified (table S1a-b). Constructs were N-terminally fused to a TwinStrep tag, C-terminally HA-tagged, and placed downstream of the *At*LORE signal peptide (AT1G61380, AA 1-21).

For *Arabidopsis thaliana mik2* CRISPR/Cas9 mutants, two target sites were selected using chopchop (https://chopchop.cbu.uib.no/, table S1c). Guide RNAs were synthesized (Twist Bioscience, USA) and stacked into a GoldenGate-adapted pUC18-based vector and assembled with FastRed-pRPS5::Cas9 in a pICSL4723 vector plant expression vector.

For *SlMIK2-A* CRISPR/Cas9 constructs, two target sites were selected using chopchop (https://chopchop.cbu.uib.no/, table S1c). Guide RNAs (gRNAs) were synthesized (Twist Bioscience, USA) and stacked with respective scaffolds in a GoldenGate-adapted vector. Stacks were ligated into a CRISPR-optimized vector with pSlUBQ10::Cas9 and kanamycin resistance marker for plant expression.

### Elicitors

Enriched *Fusarium oxysporum* elicitor (EnFOE) fractions were prepared exactly as described before (Coleman *et al*., 2021) . Concentrations refer to µg total protein per mL. Shrimp shell-derived chitin (Cat no : C9752, Sigma-Aldrich, Taufkirchen, Germany) was finely ground and extracted with ddH_2_O overnight (4°C, rotation). Concentrations refer to µg chitin powder resolved per mL water. The flg22 peptide (QRLSTGSRINSAKDDAAGLQIA) was synthesized on an Abimed EPS221 system (Abimed, Langenfeld, Germany). *At*SCOOP12 (PVRSSQSSQAGGR), *Foc*SCOOPL^FOXB_11846^ (ESSSSHSERAGGR) and *Foc*SCOOPL^F9F6H0^ (YVQGSHSSHTGGR) from *Fusarium oxysporum* f.sp. *conglutinans* (strain Fo5176) were ordered from Pepmic Corp. Ltd (China).

### Detection of ROS accumulation

Leaf discs (⌀ 4 mm) from 6-8-week-old soil-grown plants were floated on 200 µL ddH_2_O for 6 h (gain-of-function) or overnight in white 96-well plates (RT, dark). Water was replaced by 75 µL substrate containing 2 µg/mL type II horseradish peroxidase (Roche, Germany) and 5 µM L-012 (WAKO Chemicals, Germany). Luminescence was recorded as relative light units (rlu) at 1 min intervals with a Tecan F200 (Tecan, Switzerland) or HRPCS5 system (Photek, UK). Measurements included 10 min background followed by 60-120 mins after elicitor treatment. Values were normalized to mean background luminescence and mock-treated samples (included per genotype and plate). Normalized values per leaf disc were summed over the measurement period and are presented as cumulative (cum.) ROS.

### Phylogenetic analysis

Sequences of full length proteins similar to *Arabidopsis thaliana At*MIK2 of species listed in table S1d-e were acquired using BLAST (Altschul *et al*., 1990) from Phytozome (https://phytozome-next.jgi.doe.gov/; (Goodstein *et al*., 2012) or NCBI (https://blast.ncbi.nlm.nih.gov/). A search for *MIK2-clade* genes in bryophytes, lycophytes and monilophytes using the ICIPS-garden BLAST server (http://134.176.27.173/blast/; (Roessner *et al*., 2024) remained without results. Sequences were aligned using MAFFT (Katoh *et al*., 2002) . Maximum-likelihood phylogeny was generated using IQ-TREE2 (Minh *et al*., 2020) with 100 bootstraps. Computing was carried out using de.NBI VM large (28 VCPUs, 64 GB RAM). Sequences used for the analysis and gene copy numbers can be found in table S1d-f.

### Synteny analysis

A locus analysis was performed for the MIK2-clade locus of *Solanum lycopersicum* v3.2 and *Arabidopsis thaliana* ARAPORT11 as described before (Snoeck *et al*., 2024) . In short, BLASTP (BLAST 2.9.0+, e-value 10) was used to identify the syntenic loci by mining the genomes for homologues of the strongly conserved neighbor genes of *Arabidopsis AtMIK2* (AT4G08850); AT4G08840 and AT4G08920. Subsequently, putative MIK2-clade proteins were identified using BLASTP, resulting in a list of four *Sl*MIK2-clade genes. Locus comparison was performed using R (v4.0.3) and the R-package genoPlotR (v0.8.11) using the extracted contiguous MIK2-clade loci and their corresponding annotation. The resulting figure was edited in Corel-DRAW Home & Student x7.

### Aequorin luminescence measurements

7-day-old liquid-grown apoaequorin-expressing *Arabidopsis thaliana* seedlings were incubated individually in 96-well plates containing 100 µL of 5 µM coelenterazine-h solution (PJK, Germany) overnight (RT, dark). Luminescence was recorded as relative light units (rlu) using a Luminokan Ascent 2.1 (Thermo Fisher Scientific, Germany). Each well was scanned 12 times at 10 s intervals to determine background, followed by 180 measurements at 10 s intervals after elicitor treatment. Remaining aequorin was discharged by adding 150 µL discharge solution (2 M CaCl_2_, 20 % ethanol). Cytosolic calcium ion concentrations ([Ca^2+^]_cyt_) were calculated as luminescence per second relative to total luminescence remaining (*L/L_max_*) as described (Ranf *et al*., 2012) . Normalized values were summed over the measurement period and are presented as cumulative (cum.) [Ca^2+^]_cyt_.

### Ethylene measurements

For the measurement of ethylene accumulation after elicitor treatment, leaf discs (⌀ 4 mm) from youngest fully expanded leaves of 8 – 12-week-old *Solnaum lycopersicum* plants were floated on ddH_2_O overnight (RT, dark). The next day, 3 leaf discs per sample were transferred to 5 mL glass vials containing 100 µL ddH_2_O, and 50 µL elicitor or mock solution was added to indicated concentrations. Vials were sealed airtight and incubated under gentle agitation for 4.5 h. 1 mL of headspace gas was sampled with a syringe and injected into a Varian 3300 gas chromatograph (Waters Corporation, USA) equipped with a 1 m long Al_2_O_3_ column. Detector was set to 225°C, injector and column to 80°C, and ethylene was separated using O_2_, H_2_ and N_2_ at 0.5 MPa. Ethylene amounts were calculated as described (Kruedener *et al*., 1995) .

### Gene-expression analysis

For analysis of elicitor-induced gene-expression, 4-week-old hydroponically grown *Solanum lycopersicum* plants (azygous wildtype) were submerged in mock or elicitor solutions for 2 h. Whole plants were frozen in liquid nitrogen and finely ground. For *Fusarium oxysporum* f.sp*. lycopersici* strain 007 *(Fol007)*-induced gene-expression, 3 randomly chosen mock- and *Fol*007-infected plants from 5 infection experiment (fig. 4f-g) were frozen in liquid nitrogen and finely ground. For the analysis of *SlMIK2-A* splice-variants, 10 leaf discs (⌀ 4 mm) from youngest fully developed leaves of 8 – 12-week-old plants were frozen in liquid nitrogen and ground using a tissue lyzer. RNA was isolated from 3 g powder or 10 ground leaf discs using TRIzol (Roche, Switzerland) and the Direct-zol^TM^ RNA Miniprep Plus kit (Zymo Research, Germany). cDNA synthesis, including DNaseI treatment, was performed from 2 µg RNA using the ReverseAid kit with random hexamers (Cat No : K1621, Thermo Fisher Scientific, Germany). qPCR was conducted using Takyon Low ROX SYBR (Eurogentec, Belgium) with an AriaMx system (Agilent Technologies, USA). Expression was calculated using the 2^−ΔΔCT^ method (Livak & Schmittgen, 2001) and normalized to *SlEF1α* (Solyc06g009970). Mean values from two technical replicates were used. Data represents expression in treated versus mock samples. *SlMIK2-A* splice-variants are shown relative to splice-variant 1. Primers are listed in table S1b.

### Stable transformation of Arabidopsis thaliana and Solanum lycopersicum

For generation of the *A. thaliana mik2* CRISPR mutant, wildtype apoaequorin-expression Col-0^AEQ^ plants (pMAQ2) were transformed *via* floral dip using Agrobacteria (GV3101) carrying the respective constructs (see ‘Molecular cloning’). Homozygous knock-out mutation was confirmed in the T_2_ generation by PCR and sequencing, and T-DNA was segregated out using the FastRed marker. Mutation schematics are shown in fig.SIIIb.

Tomato (cv. M82) *Slmik2-A*^CRISPR^ mutants and the azygous wildtype control (construct lacking guide RNAs) were generated under sterile conditions using Agrobacteria (strain GV3101) carrying respective constructs (see ‘Molecular cloning’). Cotyledon transformation, callus formation, and regeneration were performed as described (Wittmann *et al*., 2016) . Homozygous mutations were confirmed in T_2_ generation by PCR and sequencing, and T-DNA removal by segregation was confirmed by PCR. A potential off-target (3 mismatches) of one guide RNAs (Target 2, table S1c) in the sequence of *Cf-9* (LOC101255572) was confirmed as wildtype by PCR and sequencing in all the lines.

### Agrobacteria-mediated transient transformation of *Nicotiana benthamiana*

Agrobacteria (GV3101) were transformed with desired constructs and selected on YEB plates (5 g/L beef extract, 1 g/L yeast extract, 5 g/L peptone, 5 g/L sucrose, 0.3 g/L MgSO_4_, 20 g/L agar-agar) containing antibiotics (10 µg/mL rifampicin, 30 µg/mL gentamicin, 50 µg/mL kanamycin) for 3 days at 28°C. For preculture, colonies were grown in 3 mL low-salt LB (10 g/L peptone, 5 g/L yeast extract, 5 g/L NaCl) with 30 µg/mL gentamicin and 50 µg/mL kanamycin for 24 h (28°C, 180 rpm). Cultures were inoculated (10 µL/mL) into bacterial growth medium (10 g/L peptone, 10 g/L yeast extract, 5 g/L NaCl) with 30 µg/mL gentamicin, 50 µg/mL kanamycin and 20 µM acetosyringone (Cat no : D134406, Sigma-Aldrich, Germany), and incubated for 24 h (28°C, 180 rpm). Cells were harvested (2,500xg, 5 min, RT), washed with infiltration buffer (10 mM MES pH 5.8, 10 mM MgCl_2_, 20 µM acetosyringone), resuspended to OD_600_ of 1.0, and mixed 1:1 with bacteria carrying plasmids for p19 silencing suppressor expression (Voinnet *et al*., 2003) . Suspensions were infiltrated into youngest fully-expanded leaves of 6 – 8-week-old *N. benthamiana* plants using needleless syringes and plants were returned to growth chamber.

### Immunoblotting

Proteins were separated by SDS-PAGE using an 8% resolving gel and blotted onto PVDF membranes (Amersham Hybond 0.45 µm, Cytiva, USA) using a wet-transfer system (Cat no : 1703930, Bio-Rad Laboratories, Germany). Membranes were blocked with 5% milk in PBS for 2 h, washed with PBS-T (0.05% Tween20) and incubated with primary antibodies in 10% milk in PBS-T overnight at 4°C. Membranes were washed with PBS-T (3 x 10 min) and incubated with secondary antibodies in 5% milk in PBS-T for 2 h at RT. After washing (PBS-T, 3 x 10 min), membranes were incubated with SuperSignal West Dura substrate (Thermo Fisher Scientific, Germany) and imaged using a Fusion SL Imager (Vilber Lourmat, Germany) with the accompanying software. Equal loading and transfer were confirmed by Coomassie brilliant blue or Ponceau S staining. Quantification of imaged signals was performed using the built-in feature of the Fusion SL Imager software.

### Ligand-depletion assay

Apoplastic expression of ectodomains and ligand-depletion assay were adopted from (Laohavisit *et al*., 2020; Ishihama *et al*., 2022; Shu *et al*., 2026) . *Nicotiana benthamiana* leaves were transformed *via* Agrobacteria infiltration to over-express apoplast-targeted, TwinStrep-tagged ectodomains (see ‘Molecular cloning’). 3 days post transformation, leaves were vacuum-infiltrated with cold buffer (20 mM Tris-HCl, pH 7.8), dried and rolled around 1 mL pipette tips. Tips were placed in 20 mL syringes within 50 mL tubes. Apolastic wash fluids (AWFs) were collected by centrifugation (5,000xg, 10 mins, 4°C) and stored at -20°C.

For the ligand-depletion assay, AWFs were thawed on ice, centrifuged (13,000xg, 10 mins, 4°C), and 6 mL supernatant was applied to Strep-Tactin® 4Flow® high capacity columns (Cat no : 2-1251-005, IBA Life Sciences, Germany). Columns were washed with 2 mL Buffer W and equilibrated with 1 mL BSA solution (1 mg/mL). Elicitor diluted in BSA were applied in three sequential 200 µL fractions; the final fraction was eluted with 200 µL BSA. All fractions were collected and frozen. To control for initial unspecific binding, only the third fraction (‘unbound fraction’) was used for subsequent Ca^2+^-bioassays (see ‘Aequorin luminescence measurements’). LRR ecodomains were eluted with 500 µL 1 x Buffer E, boiled in SDS-sample buffer (50 mM Tris-HCl pH 6.8, 1% β-Mercaptoethanol, 2% SDS, 0.1% bromophenol blue, 10% glycerol) and detected by immunoblotting using HA-tag specific antibodies (Cat no: 7C9, Chromotek, Germany).

### Co-Immunoprecipitation

Interactors of interest (see ‘Molecular cloning’) were co-expressed in *Nicotiana benthamiana* as described above. Agrobacteria suspensions were adjusted to OD_600_ of 1.0 in a 1:1:1 ratio (interactor 1 : interactor 2 : p19). 36 hours post infiltration, elicitor solutions were infiltrated into one half of transformed leaves using a needleless syringe, the second half received ddH_2_O as mock. After 20 min, 70 leaf discs (⌀ 4 mm) per combination were harvested in 2 mL tubes containing glass beads, snap-frozen in liquid nitrogen and finely ground using a tissue lyzer. Powder was resuspended in 700 µL extraction buffer (50 mM Tris-HCl pH 7.5, 150 mM NaCl, 10% glycerol, 0.5% NP-40, 1 mM EDTA, 1% PVPP, 1% protease inhibitor cocktail, 1 mM PMSF, 10 mM DTT), rotated at 4°C for 30 mins, diluted with 700 µL wash buffer (extraction buffer with only 0.1% NP-40 and without PVPP), centrifuged (13,000xg, 15 min, 4°C), and supernatants transferred to fresh tubes. Input samples (100 µL) were boiled in SDS-sample buffer. Remaining supernatant was added to 15 µL of GFP-Trap® Agarose (Cat no : gta, ChromoTek, Germany), which was equilibrated in wash buffer beforehand. Samples were rotated for 3 h at 4°C, supernatant removed, beads washed 4 times with 400 µL wash buffer and boiled in SDS-sample buffer. GFP- and Myc-tagged fusion proteins were detected by immunoblotting using anti-GFP and anti-Myc antibodies (Cat no : 3h9 and 9E1, ChromoTek, Germany).

### Gain-of-function assay in *Nicotiana benthamiana*

Proteins of interest (see ‘Molecular cloning’) were over-expressed in *Nicotiana benthamiana* as described above. 36 hours post infiltration, leaf discs (⌀ 4 mm) were collected for ROS measurements and protein extraction. ROS accumulation was measured 6 hours after excision as described above. For protein extraction, samples were placed in 2 mL tubes with glass beads, snap-frozen in liquid nitrogen and finely ground with a tissue lyzer. Powder was resuspended in extraction buffer (see ‘Co-Immunoprecipitation’, 20 µL/leaf disc), rotated at 4°C for 30 min, centrifuged (13,000xg, 15 min, 4°C) and supernatant boiled in SDS-sample buffer. Myc-tagged proteins were detected by immunoblotting using anti-Myc antibodies (as above).

### Confocal microscopy

To visualize subcellular localization of receptors, fluorescently labelled *At*MIK2-GFP and *Sl*MIK2-A-mCherry (see ‘Molecular Cloning’) were over-expressed in *Nicotiana benthamiana* as described above in a 1:1:1 ratio with silencing suppressor p19. Laser-scanning microscopy was performed using a Leica TCS SP5 system (Leica, Germany) with a 20x water immersion objective. GFP was excited with a 488 nm argon laser and detected 500 – 550 nm using a HyD detector. Fluorescence of mCherry was excited with a 561 nm DPSS laser and detected at 570 – 620 nm.

### Fusarium oxysporum infection assays

Root-dip method was adapted from (Mes *et al*., 1999) . *F. oxysporum* f.sp. *lycopersici* strain *Fol*007 was provided by Frank Takken, University of Amsterdam. Tomato seeds were planted in sieved soil, stratified for 7 days (dark, 4°C) and grown under long-day conditions (8h:16h, light:dark photoperiod, 20–22°C, 55% RH) for 14 days before infection. *Fol007* was grown on potato extract glucose agar (PDA) plates for 14 days (RT, 16h light). Agar-plugs from fully gown plates were transferred to sporulation medium (30 g/L sucrose, 1.7 g/L yeast nitrogen base, 10 g/L KNO_3_, autoclaved) and incubated for 4 days (dark, 28°C, 180 rpm). Spores were harvested by filtration and centrifugation, washed 3 times and resuspended to 25 mio spores/mL in sterile water. 2-week-old seedlings were uprooted, roots trimmed to 1 cm, and dipped in spore or mock (sterile water) solution for 10 min. Seedlings were repotted individually in sieved soil and grown for 3 weeks under the same conditions. For evaluation, aboveground tissue was cut at soil level. Fresh-weight and plant height (base to main stem meristem) were measured. Values of infected plants were normalized to mean values of mock-treated plants per genotype and calculated as % fresh-weight and % plant height relative to mock.

### Structural predictions

Structural alignment and visualization were performed using matchmaker in UCSF ChimeraX (University of California, USA), including models Q8VZG8 (*At*MIK2) and A0A3Q7G8D9 (*Sl*MIK2-A) fetched from UniProt and generated AlphaFold model for *Sl*MIK2-B (data repository). Complex between ectodomains of *Sl*MIK2-A (Ser21-Leu689) and *Sl*SERK3a (Asn27-Ala228) including PTMs NAG(NAG(Man(Man)(Man))) on *Sl*MIK2-A N431 and N471 was predicted using AlphaFold 3 and visualized with UCSF ChimeraX (data repository). Visualization of predicted local distance differences test (pLDDT) scores for all AF3 predictions, as well as the predicted template modeling (pTM) score and the interface predicted template modeling (ipTM) score for the *Sl*MIK2-A/*Sl*SERK3a complex can be found in the **fig.S1**.

### Statistical analysis

Statistical analyses were performed using PRISM 10.6.1 (GraphPad Software, USA) and are explained in detail in the respective figure legends.

## Results

### *Fusarium* elicitor-responsiveness and MIK2-clade proteins are conserved across plant families

Bioinformatics analyses identified SCOOP-LIKE peptide sequences embedded in proteins of *Fusarium oxysporum* f.sp. *conglutinans Fo5176* (*Foc*SCOOPL) that exhibit *At*MIK2-dependent immunogenic activity in *Arabidopsis.* We tested whether these predicted SCOOPLs share elicitor-activity with the enriched *F. oxysporum* elicitor fraction (EnFOE) in species outside the Brassicales. Similar to *At*SCOOP12, the *Foc*SCOOPL^FOXB_11846^ (Wu *et al*., 2024) and *Foc*SCOOPL^F9F6H0^ (Rhodes *et al*., 2021) peptides induced reactive oxygen species (ROS) accumulation exclusively in *A. thaliana* and *Brassica rapa*, while soybean (*Glycine max*), tomato (*Solanum lycopersicum*), and barley (*Hordeum vulgare*) showed no response (**fig. 1b-d**). In contrast, EnFOE triggered a ROS burst in most non-Brassicales species tested (fig. 1a). Notably, *Nicotiana benthamiana* did not respond to *At*SCOOP12, *Foc*SCOOPLs, or EnFOE, but responded to flg22 and chitin (**fig. 1a-d; fig.S2a-b**). These results suggest that EnFOE-induced ROS production in non-Brassicales species is mediated by immunogenic patterns distinct from the proposed *Fusarium* SCOOPLs.

**fig. 1.**
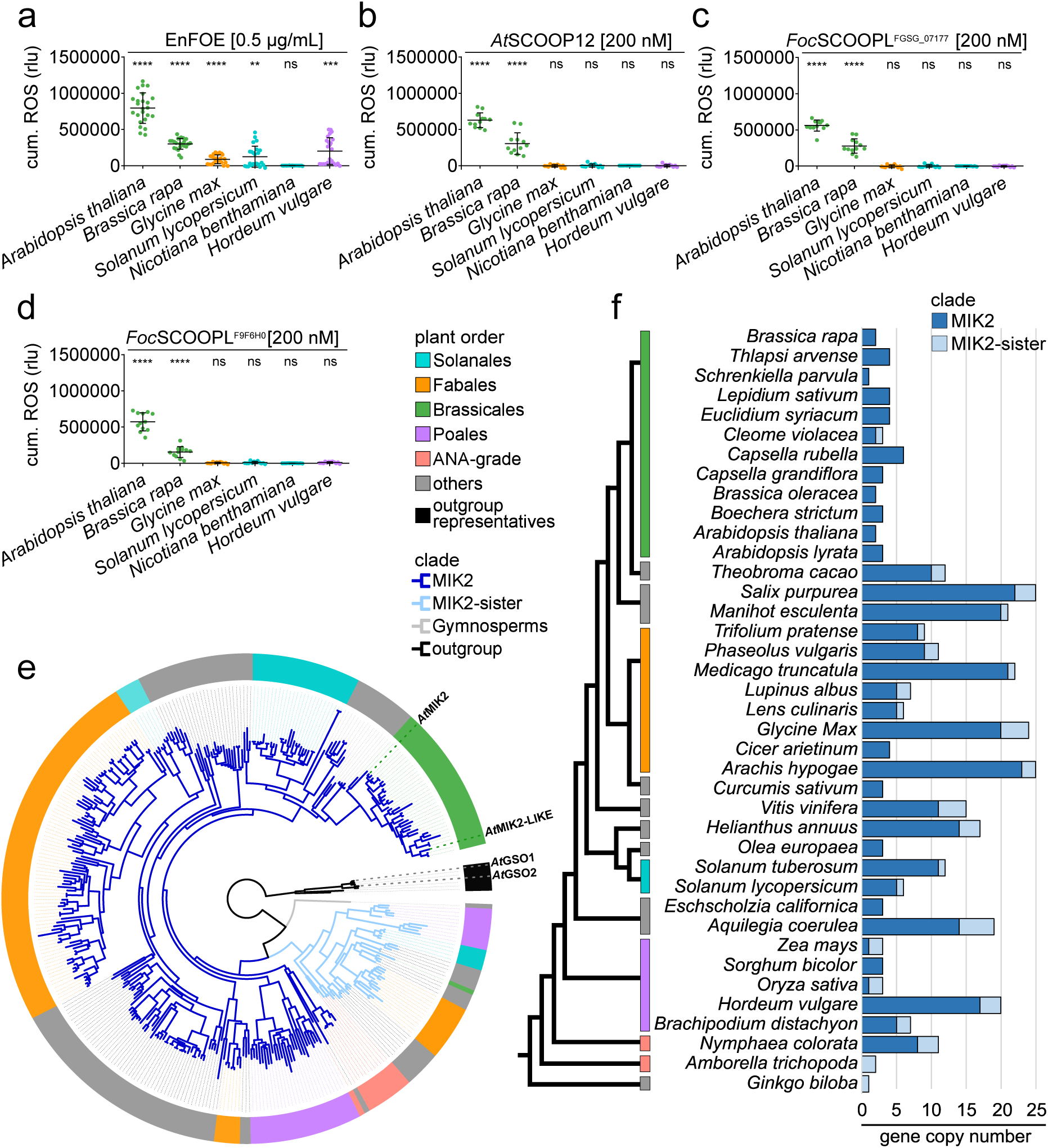
Elicitor-responsiveness and phylogeny of MIK2-clade genes in plants of different orders. (a-d) Accumulation of reactive oxygen species (ROS) measured in leaf discs from Brassicales species *Arabidopsis thaliana* and *Brassica rapa* (Pak choi), Fabales species *Glycine max* (soybean), Solanales species *Solanum lycopersicum* (tomato) and *Nicotiana benthamiana,* as well as Poales species *Hordeum vulgare* (barley) following elicitation with an enriched elicitor fraction from *Fusarium oxysporum* (EnFOE; Coleman *et al.,* 2021), 200 nM *At*SCOOP12 (Gully *et al.,* 2019), 200 nM *Foc*SCOOPL^FOXB_11846^ (Wu *et al.,* 2024) or 200 nM *Foc*SCOOPL^F9F6H0^ (Rhodes *et al.,* 2021). Recorded values were normalized to background and mock-treated samples and summed up over a period of 60 min (cumulative [cum.] ROS) and are given in relative light units (rlu). Data are pooled from 3 independent experiments, each with n = 4-12, bars indicate mean and standard deviation. Asterisks indicate statistically significant differences from respective mock-treated samples. Unpaired t-test was performed on data before normalization to mock-treated samples. **** = *P* < 0.0001, *** = *P* < 0.001, ** = *P* < 0.01, ns = not significant with *P* > 0.05. **(e)** Simplified maximum likelihood (ML) phylogeny of *MIK2* genes including representatives of major angiosperm lineages and *Ginkgo biloba* (gymnosperm). An early duplication event separated the MIK2-clade (dark blue branches) from the MIK2-sister-clade (light blue branches). Genes are color-coded according to plant orders as per legend. ANA-grade refers to basal angiosperm orders including Amborellales, Nymphaeales, and Austrobaileyales. *Arabidopsis thaliana* genes included in the analysis are specifically highlighted. A high-resolution and detailed representation of the ML phylogenetic tree can be found in fig.S2c. **(f)** Evolutionary relationship (order level) and copy number of genes belonging to the MIK2-clade (dark blue) and MIK2-sister-clade (light blue) found in the different plant species that were included in the phylogenetic analysis.

Differences in plant responsiveness to EnFOE and SCOOPs/SCOOPLs prompted us to search for *At*MIK2 orthologues in diverse plant families that may mediate *Fusarium* elicitor perception but lack SCOOP recognition. We therefore identified proteins with structural and phylogenetic similarity to *At*MIK2 outside the Brassicales. Phylogenetic analysis including representatives of major angiosperm lineages and Ginkgo revealed that MIK2-clade genes are evolutionarily conserved in seed plants (100% support) but absent in bryophytes, lycophytes, and monilophytes (**fig. 1e-f; fig.S2c**). An early duplication event in a common ancestor of monocots, eudicots and water lilies (*Nymphaeales*) gave rise to the MIK2-clade as sister to the MIK2-sister-clade (94% support), the latter uniquely containing sequences from *Amborella trichopoda* (**fig. 1e-f; fig.S2c**). Both clades include representatives from Poales, Solanales, Brassicales, and Fabales.

Within the MIK2-clade, multiple duplication events occurred, e.g., between one and two at the root of eudicots (81% and 41% support). However, most retained duplications appear at family- or species-level (**fig. 1e-f; fig.S2c**). In contrast, genes of the MIK2-sister-clade seem frequently purged after duplication and are largely absent from Brassicales (**fig. 1e-f; fig.S2c**). Conversely, MIK2-clade genes are more often retained with four (chickpea) to twenty-three (peanut) copies in Fabales, between five (tomato) and eleven (potato) copies in Solanales, and in grasses from one in rice up to seventeen copies in barley (fig. 1f**; fig.S2c**). Due to limited annotation certainty, *N. benthamiana*, *N. tabacum* and wild tomato (*Solanum chilense)* were only included in the phylogenetic analysis, precluding reliable copy number estimates (**fig. 1e-f; fig.S2c**).

Members of the Brassicales generally retain fewer MIK2-clade copies, ranging from one in *Schrenkiella parvula* and six in *Capsella rubella*, compared to other rosids and asterids such as potato (eleven copies) or sunflower (fourteen copies) (**fig. 1f; fig.S2c**). We identified five MIK2-clade genes in tomato, whereas *Arabidopsis thaliana* contains two copies – *MIK2* (AT4G08850) and *MIK2-LIKE* (AT1G35710) – arising from a Brassicales-specific duplication (100% support). In Fabales, a clade-specific duplication (100% support), followed by multiple lineage-specific duplication events, resulted in twenty retained genes in *Glycine max* (**fig. 1f; fig.S2c**). Overall, MIK2-clade genes are evolutionarily conserved in seed plants, but exhibit pronounced species-specific patterns of duplication and retention.

### A MIK2-clade protein from *Solanum lycopersicum* shows similar properties to *At*MIK2 in *Fusarium* elicitor perception

To further investigate putative *At*MIK2 orthologues involved in *Fusarium* elicitor perception, we focused on MIK2-clade proteins from *S. lycopersicum*, a species frequently affected by *Fusarium* infections (McGovern, 2015) . We cloned all five tomato MIK2-clade genes and the MIK2-sister-clade gene, designated *SlMIK2-A to -E* and *SlMIK2-SISTER*, respectively (**fig. 2a**), and transiently expressed them in *N. benthamiana*, which is naturally unresponsive to EnFOE- or SCOOP-treatment (**fig. 1a-d**). Wildtype *At*MIK2 and a kinase-dead variant (*At*MIK2^km, K802A) were included as controls, as this system has been validated previously (Coleman *et al*., 2021; Rhodes *et al*., 2021) . All constructs were expressed, with *Sl*MIK2-E showing lower accumulation (**fig.S3a**).

**fig. 2.**
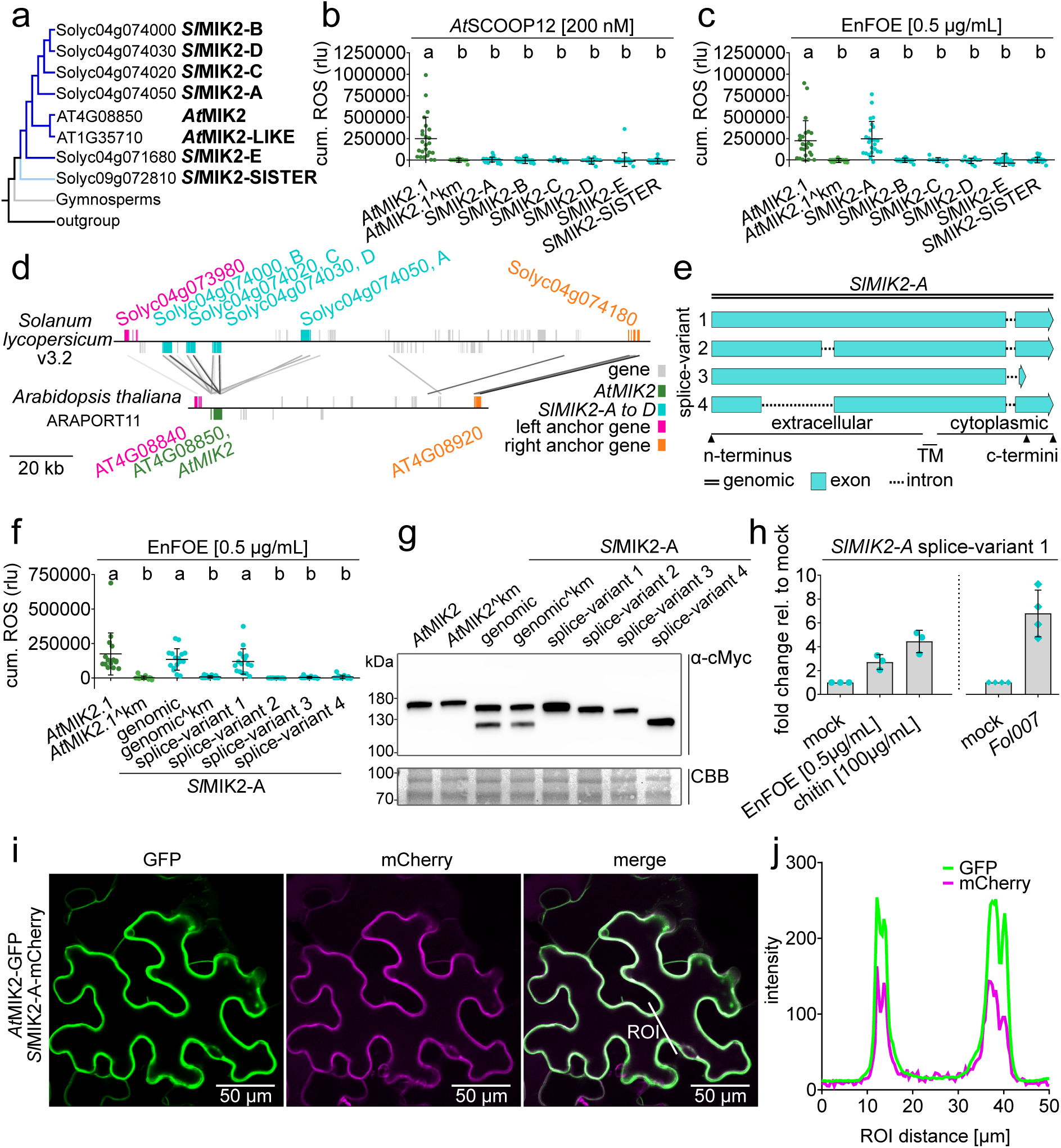
Analysis of MIK2-clade proteins from *Solanum lycopersicum*. **(a)** Simplified phylogenetic tree of MIK2- and MIK2-sister-clade proteins from *Arabidopsis thaliana* and *Solanum lycopersicum* with gene identifiers and nomenclature used in this study**. (b-c)** EnFOE- and *At*SCOOP12-induced accumulation of reactive oxygen species (ROS) measured in leaf discs from *Nicotiana benthamiana* transiently expressing *Arabidopsis thaliana* MIK2 (*At*MIK2), a kinase-inactive version of *At*MIK2 (*At*MIK2^km, K802A) or *Solanum lycopersicum* proteins from the MIK2- and MIK2-sister-clades (*Sl*MIK2-A to -E, *Sl*MIK2-SISTER). Recorded values were normalized to background and mock-treated samples and summed over a period of 60 min (cumulative [cum.] ROS) and are given in relative light units (rlu). Data are pooled from 4-7 independent experiments, each with n = 4, bars indicate mean and standard deviation. Letters show significant differences (one-way ANOVA, Tukey’s test, *P* < 0.0003). **(d)** Comparison of the contiguous *MIK2* locus in *Arabidopsis thaliana* and *Solanum lycopersicum.* Anchor genes, *SlMIK2-A* to *-D, AtMIK2*, and other genes are colored as per legend. BLAST hits are indicated with lines (e-value < 1e-02) with scores according to grayscale gradient with darker grays indicating higher similarity. Gene orientation is depicted by the position above or below the line. Size of the genomic regions can be compared on the bottom left and is given in 20 kilobases (kb) **(e)** Graphical representation of predicted splice-variants 1 to 4 from *SlMIK2- A.* **(f)** EnFOE-induced accumulation of reactive oxygen species (ROS) measured in leaf discs from *Nicotiana benthamiana* transiently over-expressing *At*MIK2, *At*MIK2^km and different versions of *Sl*MIK2-A (genomic sequence, K791A mutated ^km version and coding sequence of individual splice-variants). Recorded values were normalized to background and mock-treated samples and summed up over a period of 60 min (cum. ROS) and are given in relative light units (rlu). Data are pooled from 4 independent experiments, each with n = 4, bars indicate mean and standard deviation. Letters show significant differences (one-way ANOVA, Tukey’s test, *P* < 0.0001). **(g)** Western blot visualization of C-terminally cMyc-tagged proteins corresponding to data of one experiment from fig.2f and fig.SIId. Blots were probed with α-cMyc antibodies. Approximate protein size is marked on the left edge and given in kilodaltons (kDa). Equal protein loading was confirmed by Coomassie Brilliant Blue staining (CBB). **(h)** Induction of *SlMIK2-A* (splice-variant 1) transcripts in whole hydroponically grown *Solanum lycopersicum* cv. M82 wildtype plants, 2 hours after EnFOE and chitin treatment (left) or in shoot stems of soil-grown plants 21 days after infection with *Fusarium oxysporum* f.sp. *lycopersici* strain *Fol007* (right). Values were normalized to the expression of the housekeeping gene *SlEF1α* and are shown as fold-change relative to respective mock-treated controls. Individual symbols show results from 3 (elicitor-treatment, dots) or 4 (*Fol007* infection, squares) biological replicates **(i)** Confocal microscopy images of GFP- (left) and mCherry- (middle) fluorescence detected in *Nicotiana benthamiana* leaves transiently co-expressing *At*MIK2-GFP and *Sl*MIK2-A-mCherry (splice-variant 1). Merge of GFP and mCherry channels is shown on the right. White line in the merge image indicates the ROI (region of interest) used to calculate fluorescence intensity shown in fig.2j. **(j)** GFP and mCherry fluorescence intensity in arbitrary units along the ROI line indicated in the merge image in fig.2i.

As expected, *At*SCOOP12 induced a robust ROS burst in leaf discs expressing *At*MIK2, but not in those expressing *At*MIK2^km or either of the tomato proteins (**fig. 2b**). Similarly, *Foc*SCOOPLs triggered ROS production exclusively in leaves expressing wildtype *At*MIK2 (**fig.S3b-c**). EnFOE induced ROS accumulation in leaves expressing wildtype *At*MIK2, and likewise, the expression of *Sl*MIK2-A conferred EnFOE-sensitivity to *N. benthamiana,* whereas all other tomato MIK2-related proteins did not (**fig. 2c**). These results indicate that *Sl*MIK2-A may function similarly to *At*MIK2 in the perception of EnFOE, but does not mediate responses to *At*SCOOPs or *Foc*SCOOPLs.

Consistent with a shared evolutionary origin, synteny analysis revealed a conserved genomic region containing *At*MIK2 and four tomato MIK2-clade genes in the respective species (**fig. 2d**). *SlMIK2-B* to -*D* are arranged in tandem on the minus strand, while *SlMIK2-A* resides on the plus strand and is separated by an unrelated gene (**fig. 2d**).

*SlMIK2-A* encodes a 1048-amino acid protein predicted to be an LRR-RK containing 25 LRRs and shares 48% sequence identity and 65% similarity with *At*MIK2 (1045 amino acids, 24 LRRs). Notably, *SlMIK2-A* is predicted to generate four splice-variants (**fig. 2e**), which likely explains the characteristic double-band observed on western blots, as the C-terminal cMyc-tag enables detection of the three variants sharing an identical C-terminus (**fig. 2e, 2g; fig.S3a**). Although splice-variant 1 is most similar to *At*MIK2, all four splice-variants were cloned individually according to the respective coding sequences and by mutation of additional splice-sites.

Each splice-variant was transiently overexpressed in *N. benthamiana*, alongside the unmodified (genomic) and a kinase-dead version of *Sl*MIK2-A (^km, K791A), as well as *At*MIK2 controls (**fig. 2g**). Only splice-variant 1 conferred EnFOE-sensitivity comparable to *At*MIK2 and genomic *Sl*MIK2-A (**fig. 2f**), whereas none of the variants conferred *At*SCOOP12-sensitivity (**fig.S3d**). As observed for *At*MIK2, *Sl*MIK2-A requires a functional kinase domain to confer EnFOE-sensitivity (**fig. 2f**). RT-qPCR analysis using variant-specific primers showed that splice-variant 1 is the predominant transcript in wildtype tomatoes, while splice-variants 2 and 4 were 74% and 55% less abundant, respectively, and splice-variant 3 was barely detectable (**fig.S3e**). In summary, the predominant splice-variant 1 of *SlMIK2- A* is the only one translated into a protein that can confer EnFOE-sensitivity to *N. benthamiana*. The expression of this splice-variant was also transcriptionally induced 2 hours after EnFOE or chitin treatment in whole plants and 21 days after infection with *F. oxysporum* f.sp. *lycopersici* (strain *Fol007*) in shoot stems (**fig. 2h**).

Similar to *At*MIK2 (van der Does *et al.,* 2017), also *Sl*MIK2-A (splice-variant 1) localizes to the plasma membrane. Co-expression of *At*MIK2-GFP and *Sl*MIK2-A-mCherry in *N. benthamiana* followed by confocal imaging revealed overlapping fluorescence signals at the cell periphery, as confirmed by intensity profiling along the region of interest (ROI) (**fig. 2i-j**).

### *Sl*MIK2-A forms ligand-induced complexes with tomato BAK1 orthologues and depletes elicitor-activity from EnFOE

Since *At*MIK2 forms ligand-induced complexes with the co-receptor *At*BAK1 after treatment with EnFOE, *At*SCOOPs and *Fusarium* SCOOPLs (Coleman *et al*., 2021; Hou *et al*., 2021; Rhodes *et al*., 2021), we examined whether analogous complexes form between the corresponding tomato proteins. Tomato encodes two *At*BAK1 orthologues, termed *Sl*SERK3a and *Sl*SERK3b, both implicated in immunity (Mantelin *et al*., 2011; Peng & Kaloshian, 2014) . We therefore tested ligand-induced interactions between *Sl*MIK2-A and each BAK1 orthologue by co-immunoprecipitation (CoIP) after transient expression in *N. benthamiana*. The receptor pair from *Arabidopsis* was used as a positive control, while *Sl*MIK2-B, which does not mediate EnFOE perception (**fig. 2c**), was included as a negative control.

Consistent with the gain-of-function assays (**fig. 2b; fig.S3c**), *At*SCOOP12 and the *Foc*SCOOPL^FOXB_11846^ peptide induced complex formation between *At*MIK2 and *At*BAK1, whereas no interactions involving the corresponding tomato proteins were detected (**fig. 3b; fig.S4a**). In contrast, EnFOE induced complex formation between *At*MIK2 and *At*BAK1 but also between *Sl*MIK2-A and either *Sl*SERK3a or *Sl*SERK3b (**fig. 3a**). However, EnFOE did not promote association between *Sl*MIK2-B and either *Sl*SERK proteins (**fig. 3a**). Together, these results indicate similarities in the perception mechanisms of the *Fusarium* elicitor in *Arabidopsis* and tomato.

**fig. 3.**
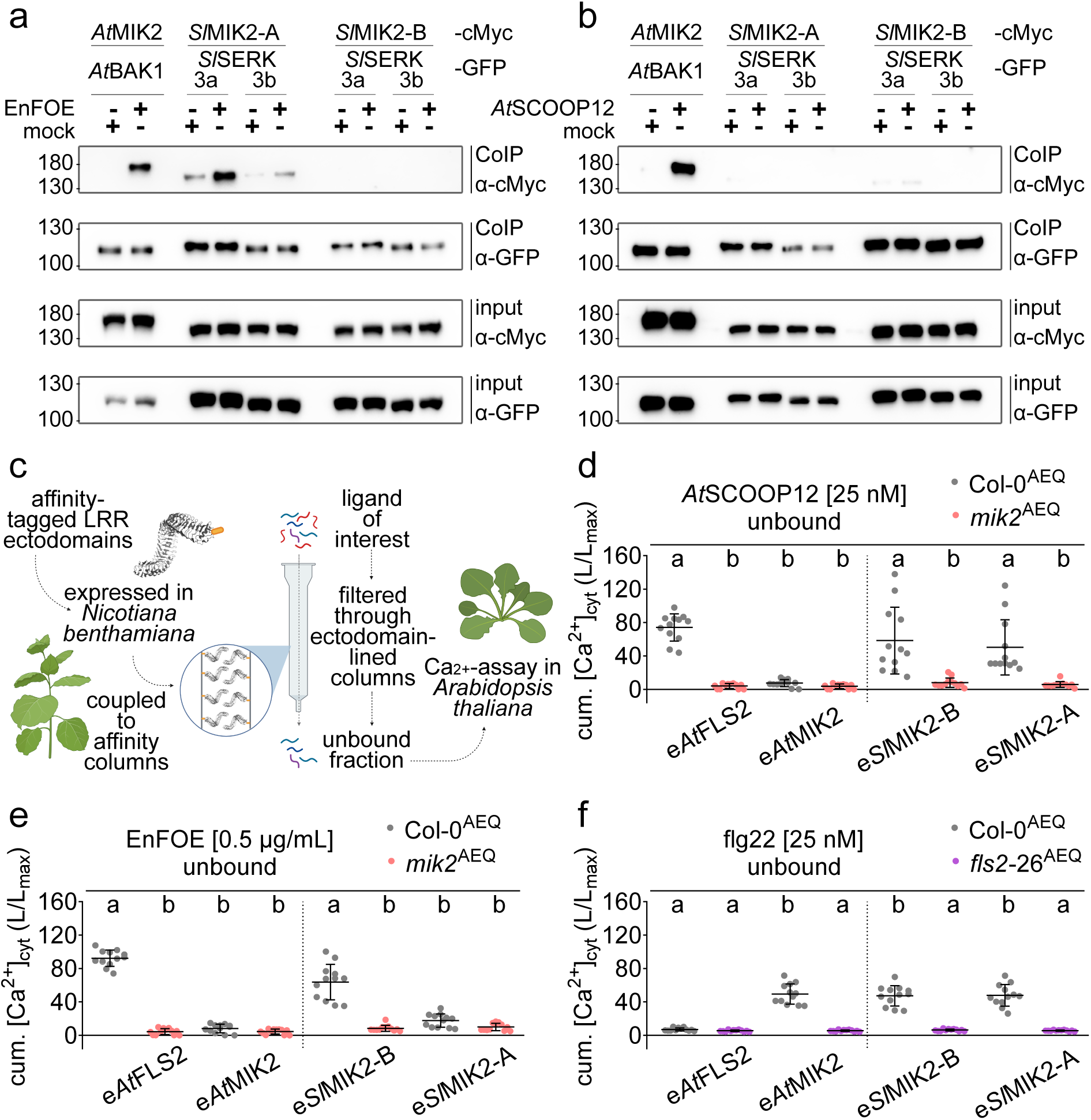
Ligand-induced receptor-complex formation and ligand-depletion assay. (a-b) Co-immunoprecipitation of MIK2 and BAK1 proteins from *Arabidopsis thaliana* (*At*MIK2-cMyc, *At*BAK1-GFP) and respective orthologues of *Solanum lycopersicum* (*Sl*MIK2-A-cMyc [splice-variant 1], *Sl*MIK2-B-cMyc, *Sl*SERK3a- or *Sl*SERK3b-GFP) transiently overexpressed in *Nicotiana benthamiana*, 20 min after elicitation with water (mock), an enriched elicitor fraction from *Fusarium oxysporum* (EnFOE) or *At*SCOOP12 (1 µM). Immunoprecipitation was performed using magnetic anti-GFP beads, western blots were probed with α-cMyc or α-GFP antibodies. Approximate protein sizes are labelled on the left and given in kilodaltons (kDa). **(c)** Workflow for the ligand-depletion assay. TwinStrep affinity-tagged ectodomains of leucine-rich repeat receptor kinases (LRR-RKs) are transiently overexpressed in *Nicotiana benthamiana*, collected with apoplastic wash fluids and coupled to Strep-Tactin gravity-flow columns. Ligands of interest are filtered through the ectodomain-lined columns and resulting unbound fractions are subsequently analyzed for remaining elicitor-activity in a Ca^2+-^assay using apoaequorin-expressing *Arabidopsis thaliana* seedlings. This figure was created in BioRender (https://BioRender.com/5xkijxt). **(d-f**) Cumulative cytosolic elevation of calcium ion concentration (cum. [Ca^2+^]cyt) in *Arabidopsis thaliana* Col-0^AEQ^ wildtype, *mik2*^AEQ^ and *fls2*-26^AEQ^ mutant seedlings after elicitation with *At*SCOOP12 (25 nM) unbound, EnFOE [0.5 µg/mL] unbound or flg22 (25 nM) unbound. Unbound fractions resulted from filtering of respective ligands through columns lined with the ectodomains of *Arabidopsis thaliana* MIK2 (e*At*MIK2) and FLS2 (e*At*FLS2) or *Solanum lycopersicum* MIK2-A [splice-variant 1] and MIK2-B (e*Sl*MIK2-A/B). Data shown are pooled from three independent experiments, each with n = 4. Values of individual seedlings (each plotted dot) were summed up over a period of 30 min after elicitation. Relative calcium levels are presented as luminescence counts per second relative to total luminescence counts remaining (L/Lmax). Bars indicate mean and standard deviation. Letters indicate significant differences within individual data sets that are separated by the dotted line (one-way ANOVA, Tukey’s test, *P* < 0.001).

**fig. 4.**
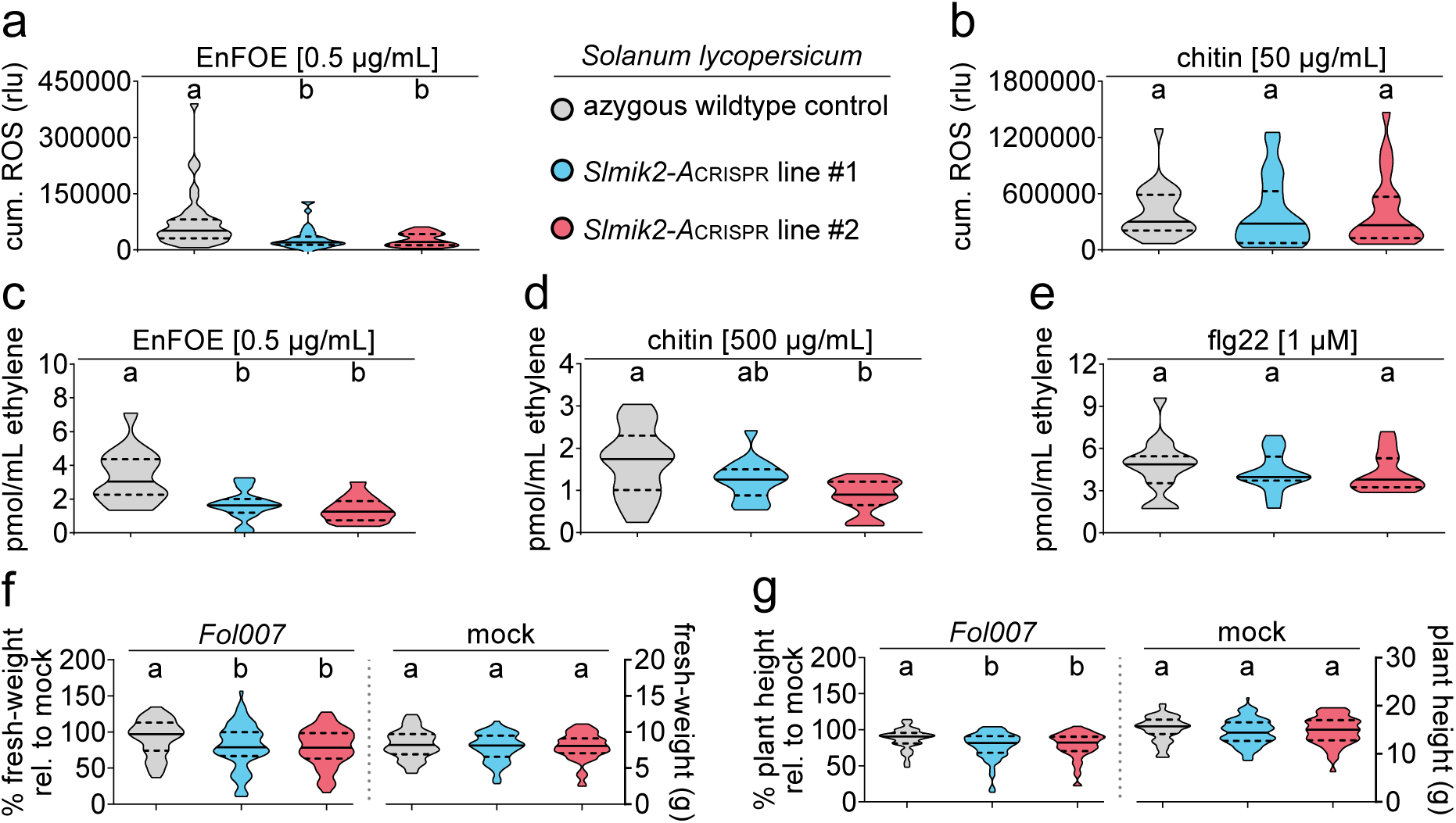
Elicitor-induced PTI responses and *Fusarium oxysporum* susceptibility of tomato *Slmik2- A* mutants. (a-b) Accumulation of reactive oxygen species (ROS) measured in leaf discs of two independent *Solanum lycopersicum Slmik2-A* CRISPR knockout mutants (line #1 blue triangles, line #2 red squares) and the corresponding azygous wildtype control (gray dots) after elicitation with an enriched elicitor fraction from *Fusarium oxysporum* (EnFOE; 0.5 µg/mL) or chitin (50 µg/mL). Recorded values were normalized to mock-treated samples and summed over a period of 120 min (cum. ROS) and are given in relative light units (rlu). Data are pooled from 3 independent experiments, each with n = 8-16 (total n = 22-36), bars indicate mean and standard deviation. Letters show significant differences (one-way ANOVA, Tukey’s test, *P* < 0.0004). **(c-e)** Ethylene concentration emitted from leaf discs of genotypes stated in a-b, 3.5 h after treatment with EnFOE (0.5 µg/mL), chitin (500 µg/mL) or flg22 (1 µM). Values were normalized to mock- treated samples. Data are pooled from 4 independent experiments, each with n = 4 (total n = 16), bars indicate mean and standard deviation. Letters indicate significant differences (one-way ANOVA, Tukey’s test, *P* < 0.001). **(f-g)** Relative changes in fresh-weight and plant height of indicated tomato genotypes caused by 21 days of infection with *Fusarium oxysporum* f.sp. *lycopersici* (strain *Fol007*). Values were calculated relative to the average of respective mock-treated plants (100%) (left panels). Fresh-weight (in g) and plant height (in cm) of mock-treated plants included in the experiments are shown in the right panels. Data are pooled from 6 independent experiments, each with n = 5-15 (total n = 57-69). Violin plots show median (solid black line) and interquartile range (area between the dashed black lines). Letters indicate significant differences within each panel (one-way ANOVA, Tukey’s test, *P* < 0.03).

Since the CoIP experiments suggest that one or several elicitor-active molecules present in EnFOE interact with the ectodomains of *At*MIK2 and *Sl*MIK2-A, we sought to further substantiate this supposition by adopting a ligand-depletion assay (Shu *et al*., 2026) (**fig. 3c**). Affinity-tagged ectodomains (N-terminal TwinStrep) of selected LRR-RKs were expressed *in planta* and collected with apoplastic wash fluids (AWFs). The ectodomains were then immobilized on affinity columns, while other apoplastic molecules were washed out. Preparations containing elicitor-activity were run through the ectodomain-lined columns and the flow-through was analyzed for depletion of the ligands. Because all tested ligands induce a transient elevation of cytosolic-free calcium concentrations ([Ca^2+^]_cyt_), remaining elicitor-activity was quantified using a sensitive calcium bioassay in apoaequorin-expressing *A. thaliana* Columbia-0 wildtype (Col-0^AEQ^) and a *mik2*^AEQ^ knockout line we generated by CRISPR/Cas9 in the same background (**fig.S4b**).

The ectodomains of *At*MIK2 (e*At*MIK2) and *Sl*MIK2-A (e*Sl*MIK2-A) were robustly expressed in *N. benthamiana*. As controls, we additionally expressed the ectodomains of *A. thaliana* FLAGELLIN-SENSING 2 (e*At*FLS2) and *Sl*MIK2-B (e*Sl*MIK2-B), which are not expected to interact with our ligands of interest (**fig.S4c**). As anticipated, columns lined with e*At*MIK2 depleted elicitor-activity from *At*SCOOP12 preparations that consequently did not trigger calcium transients that differed between Col-0^AEQ^ wildtype and *mik2*^AEQ^ mutant seedlings (**fig. 3d**). In contrast, *At*SCOOP12 preparations passed through columns lined with e*At*FLS2 elicited a clear *At*MIK2-dependent calcium response in Col-0^AEQ^ seedlings (**fig. 3d**). Similarly, neither e*Sl*MIK2-A nor e*Sl*MIK2-B depleted *At*SCOOP12, as both flow-throughs retained full elicitor-activity (**fig. 3d**). Thus, *At*SCOOP12 was specifically depleted by ectodomains of its receptor *At*MIK2.

We then applied the same setup for EnFOE preparations. Similar to *At*SCOOP12, columns lined with e*At*MIK2 fully depleted EnFOE of elicitor-activity, whereas e*At*FLS2-lined columns did not (**fig. 3e**). Notably, e*Sl*MIK2-A also depleted the elicitor-activity, as the corresponding flow-through failed to induce a calcium response in wildtype seedlings that significantly exceeded that of the *mik2*^AEQ^ mutants. Preparations filtered through e*Sl*MIK2-B-lined columns still harbored *At*MIK2-dependent elicitor-activity (**fig. 3e**). The ectodomain of tomato *Sl*MIK2-A could thus specifically deplete elicitor-activity from EnFOE that is perceived in an *At*MIK2-dependent manner in *A. thaliana*. This suggests that ectodomains of *At*MIK2 and *Sl*MIK2-A have an affinity to a similar elicitor-activity in EnFOE.

Additionally, we performed the assay using the flg22 peptide in which only flowthroughs from columns lined with ectodomains of its cognate receptor *At*FLS2 were depleted of elicitor-activity, resulting in no detectable calcium transient in wildtype seedlings (**fig. 3f**). Comparable loading of ectodomains was confirmed by western blot analysis of proteins eluted from the columns. Representative blots of one experiment for each pair can be found in **fig.S4c**, signal quantification of all three replicates can be found in **fig.S4d-i**.

Together, these results further support a dual role of *At*MIK2 in perceiving both endogenous SCOOP peptides and an exogenous *Fusarium* elicitor, whereas the phylogenetically related *Sl*MIK2-A selectively interacts with the EnFOE-associated elicitor-activity and cannot function in SCOOP perception.

### *Sl*MIK2-A is important for EnFOE-induced PTI responses and resistance against *Fusarium oxysporum* infection in tomato

To validate the role of *Sl*MIK2-A in EnFOE-induced PTI responses in its native context, we generated two independent CRISPR/Cas9 knockout lines in tomato. Both *Slmik2-A*^CRISPR^ mutants carry homozygous frameshift mutations within the first 500 bp of the coding sequence, introducing premature stop codons that disrupt all 4 splice-variants (**fig.S4j**). The corresponding wildtype control was transformed with a construct lacking the guide RNAs and has undergone tissue culture alongside the CRISPR mutants (azygous wildtype control). T-DNAs were segregated out in subsequent generations.

We first assessed elicitor-induced ROS production in the tomato mutants. Upon EnFOE treatment, both *Slmik2- A*^CRISPR^ lines showed detectable but significantly reduced ROS accumulation when compared to the azygous wildtype control (**fig. 4a**). In contrast, wildtype and mutants responded similarly to the fungal MAMP chitin (**fig. 4b**). Likewise, EnFOE-induced ethylene accumulation was significantly reduced in both mutants (**fig. 4c**). Interestingly, chitin-induced ethylene production was also significantly reduced in *Slmik2- A*^CRISPR^ line #2. Notably, even wildtype plants required relatively high concentrations of chitin (500 µg/mL) for the induction of a measurable accumulation of ethylene (**fig. 4d**). By contrast, flg22-induced ethylene production was similar across all genotypes (**fig. 4e**), indicating that loss of *Sl*MIK2-A does not cause a general defect in MAMP-induced ethylene accumulation. Taken together, similar to *At*MIK2 in *A. thaliana*, *Sl*MIK2-A is required for full responsiveness to EnFOE in tomato. We next examined whether *Sl*MIK2-A contributes to resistance to *F. oxysporum* f.sp. *lycopersici*. We inoculated two-week-old seedlings with the isolate *Fol007* via root-dip infection (Mes *et al*., 1999), assessed growth parameters three weeks post-infection and calculated percentages relative to mock-treated plants, to compare the negative effects of *Fol007* infection between the genotypes. Infected azygous wildtype controls exhibited an average fresh-weight reduction of around 8%, whereas both *Slmik2- A*^CRISPR^ lines showed significantly greater reductions of around 21% (**fig. 4f**, left). Similarly, plant height was reduced by about 12% in wildtype plants but by approximately 21% in the mutants (**fig. 4g**, left). Importantly, in mock-treated plants, fresh-weight and plant height did not differ between wildtype and mutant plants (**fig. 4f-g**, right). Thus, loss of *Sl*MIK2-A does not affect basal growth but markedly increases susceptibility of tomato plants to *Fusarium* infection.

### Conserved residues on the ectodomains of *At*MIK2 and *Sl*MIK2-A are important for conferring EnFOE-sensitivity to *Nicotiana benthamiana*

In contrast to EnFOE, interactions between SCOOPs and *At*MIK2 have already been extensively characterized. Two binding pockets engage the conserved SCOOP SxS motif *via* hydrogen bonds, with *At*MIK2 residues D246, N268, S292, H294, and H316 playing central roles (Jia *et al*., 2024; Snoeck *et al*., 2024; Wu *et al*., 2024) . Interestingly, AlphaFold-based structural alignment revealed that these critical residues are not conserved on *Sl*MIK2-A (**fig. 5a**). To determine the role of the SCOOP-binding residues of *At*MIK2 in the recognition of EnFOE, we generated *At*MIK2^D246G/N268A^ and *At*MIK2^S292A/H316A^ mutants as well as the respective quadruple mutant and transiently expressed them alongside *At*MIK2^wildtype^ in *N. benthamiana*. As previously reported, the ability of the mutants to confer *At*SCOOP12-sensitivity was abolished or significantly reduced (**fig. 5b; fig.S5k**). Similarly, EnFOE-induced ROS responses were drastically reduced for all tested mutants (**fig. 5c; fig.S5k**). Conversely, substitution mutants of the corresponding structural equivalents on the tomato receptor were still able to confer EnFOE-sensitivity in a comparable manner as the wildtype protein (**fig. 5d; fig.S5l**).

**fig. 5.**
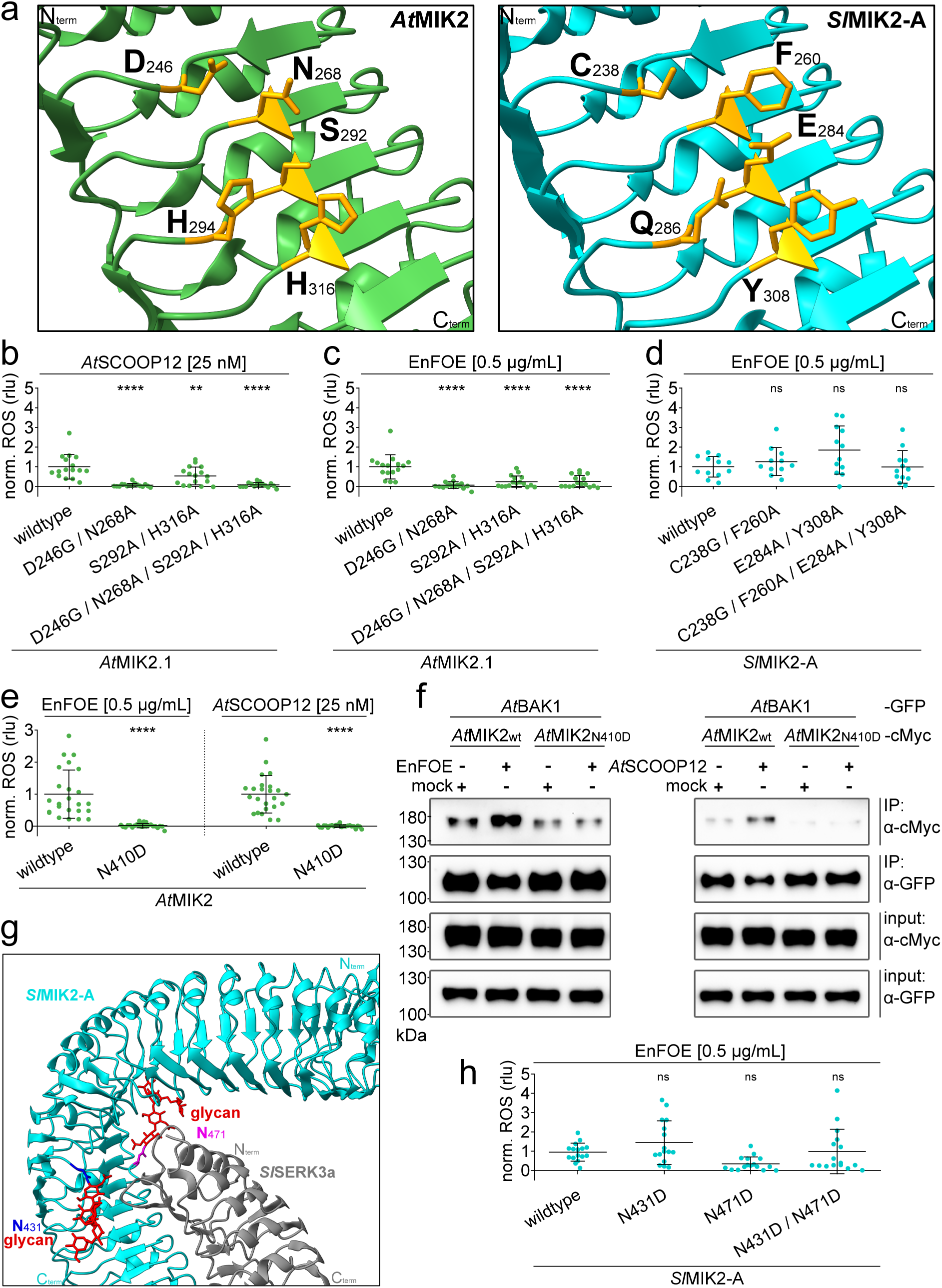
Influence of SCOOP-binding residues and *N*-glycosylation sites on EnFOE-sensitivity conferred by *At*MIK2 and *Sl*MIK2-A. **(a)** Close-up of AlphaFold (AF) structures of *At*MIK2 (left, green) and *Sl*MIK2-A (right, turquoise) ectodomains. Important SCOOP-binding residues (Jia *et al.,* 2024; Snoeck *et al.,* 2024; Wu *et al.,* 2024) on *At*MIK2 and their structural equivalents (same amino acid positions according to structural alignment) on *Sl*MIK2-A are highlighted in orange and respective amino acids and positions are indicated. Orientations of N-termini (N^term^) and C-termini (C^term^) are shown in the corners. **(b-e, h)** EnFOE- or *At*SCOOP12-induced accumulation of reactive oxygen species (ROS) measured in leaf discs from *Nicotiana benthamiana* transiently overexpressing *Arabidopsis thaliana* MIK2 (*At*MIK2.1) or *Solanum lycopersicum* MIK2-A (*Sl*MIK2-A). Proteins were expressed as wildtype or with indicated amino acid substitutions. Recorded values were normalized to background and mock-treated samples and summed over a period of 60 min (cumulative [cum.] ROS) and are given in relative light units (rlu). To enhance comparability, all individual values were standardized to average values of wildtype controls within each independent experiment (collective values of wildtype average at 1.0). Data are pooled from 3-6 independent experiments, each with n = 4, bars indicate mean and standard deviation. Asterisks indicate statistically significant differences compared to respective wildtype with **** = *P* < 0.0001, ** = *P* < 0.01, ns = not significant with ρ > 0.05, ordinary one-way ANOVA with Dunnett’s multiple comparisons test (b-d, h) or unpaired t-test (e). **(f)** Co-immunoprecipitation of *Arabidopsis thaliana* BAK1 (*At*BAK1-GFP), MIK2 wildtype (*At*MIK2^wt^-cMyc) and MIK2 with N410D substitution (*At*MIK2^N410D^-cMyc) transiently overexpressed in *Nicotiana benthamiana*, 20 min after elicitation with water (mock), EnFOE [0.5 µg/mL], or *At*SCOOP12 [25 nM]. Immunoprecipitation (IP) was performed using magnetic anti-GFP beads, Western blots were probed with α-cMyc or α-GFP antibodies. Approximate protein sizes are labelled on the left and given in kilodaltons (kDa). **(g)** AlphaFold-Multimer (AFM)-predicted complex structure of *Solanum lycopersicum* MIK2-A (*Sl*MIK2-A, turquoise) and SERK3a (*Sl*SERK3a, gray) ectodomains. Positions of putative *N*-glycosylation sites on *Sl*MIK2-A (N431, dark blue; N471, lilac) are highlighted and exemplary depicted glycan chains are shown in red. Orientation of N-termini (N^term^) and C-termini (C^term^) are shown in the corners and colored to match the respective proteins. **(h)** see description b-e.

In addition to SCOOP-binding residues, glycans *N*-linked to *At*MIK2^N410^ mediate receptor activation by enabling interaction with the co-receptor BAK1 (Jia *et al*., 2024; Wu *et al*., 2024) . To test whether *N*-glycosylation is also required for EnFOE-induced responses, we analyzed an *At*MIK2^N410D^ mutant in *N. benthamiana*. Indeed, the mutation abolished both EnFOE- and *At*SCOOP12-induced ROS responses (**fig. 5e; fig.S5h**). Co-immunoprecipitation in transiently transformed *N. benthamiana* further showed that the ligand-induced enhancement of *At*MIK2-*At*BAK1 interaction is lost in *At*MIK2^N410D^ after *At*SCOOP12 and EnFOE treatment (**fig. 5f**).

AlphaFold-based structural alignment of *At*MIK2 and *Sl*MIK2-A showed that N410 is not conserved on *Sl*MIK2-A. To identify alternative *N*-glycosylation sites potentially involved in EnFOE responses, we predicted a *Sl*MIK2-A-*Sl*SERK3a complex using AlphaFold-Multimer and identified *Sl*MIK2-A^N431^ and *Sl*MIK2-A^N471^ as candidates (**fig. 5g**). However, mutating these residues did not significantly reduce the ability to confer EnFOE-sensitivity to *N. benthamiana* (**fig. 5h; fig.S5j**).

We next sought to identify shared amino acids required for EnFOE-induced responses by both receptors. Guided by our previous analysis, we focused on surface-exposed residues conserved between *At*MIK2 and *Sl*MIK2-A but not *Sl*MIK2-B, located near alternatively spliced regions that generate non-functional *Sl*MIK2-A variants, and positioned close to or aligned with SCOOP-binding residues. By applying these criteria, we identified Y318, K345, D367, Q369 on *At*MIK2 that align with Y310, K337, D359, Q361 on *Sl*MIK2-A as candidates (**fig. 6a; fig.S6a**). We again cloned alanine-substitution mutants of all respective residues and tested them in *N. benthamiana*. Except for *At*MIK2^K345A^, all tested single mutants showed a significantly reduced ability to confer EnFOE-sensitivity (**fig. 6b, 6d**). In contrast, neither of the *At*MIK2 single alanine substitutions had a significant effect on *At*SCOOP12-induced ROS responses (**fig.S5a; fig.S6b**). While the expression levels of individual proteins could vary slightly within individual experiments (**fig.S5**), signal quantification of blots from all biological replicates show no significant differences between detected signals between any of the proteins that were compared **(fig.S7**), indicating that observed changes in ROS responses occur without corresponding changes in protein expression levels. When testing double mutants of all combinations, reductions in EnFOE-induced responses became more pronounced for both receptors and were nearly abolished in all *At*MIK2 mutants containing the Y318A substitution as well as in the *Sl*MIK2-A^D359A/Q361A^ mutant (**fig. 6c, 6e; fig.S5c-d**). For *At*SCOOP12-induced ROS responses, reductions were only significant for the *At*MIK2^Y318A/D367A^ double mutant (**fig.S5c; fig.S6c**). The observed additive effect continued in the triple mutants, where all *At*MIK2 mutants including the Y318A substitution could not confer EnFOE-sensitivity, while the effect in *Sl*MIK2-A was strongest when the other three residues were substituted by alanine (**fig.S5e-f; fig.S6d, S6f**). For *At*SCOOP12, all *At*MIK2 triple mutants containing the Y318A substitution showed a significant reduction, while sensitivity conferred by AtMIK2^K345A/D367A/Q369A^ was still comparable to that of the wildtype protein, which is in line with the proposed role of Y318 in the interaction with the C-terminal part of the SCOOP peptide (**fig.S5e; fig.S6e**; Jia *et al.,* 2024). Accordingly, the tested quadruple mutants also could not confer EnFOE- or *At*SCOOP12-sensitivity (**fig.S5g, S5i; fig.S6g**).

**fig. 6.**
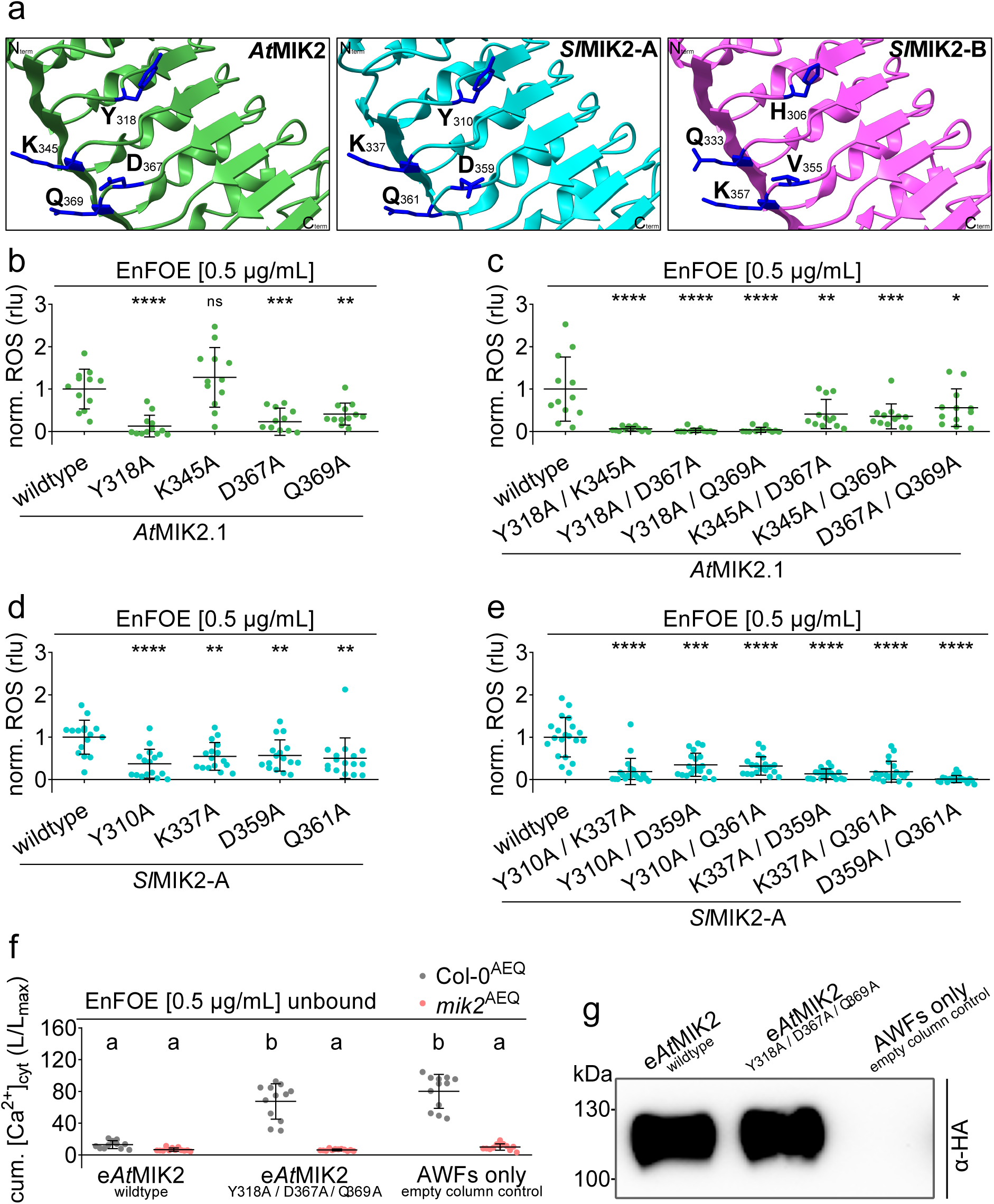
Function of conserved residues for EnFOE-sensitivity and ligand-depleting activity conferred by *At*MIK2 and *Sl*MIK2-A. **(a)** Close-up of AlphaFold (AF) structures of *At*MIK2 (left, green), *Sl*MIK2-A (middle, turquoise), and *Sl*MIK2-B (right, pink) ectodomains. Aligned surface-exposed residues conserved between *At*MIK2 and *Sl*MIK2-A, but not *Sl*MIK2-B, are highlighted in dark blue and respective amino acids and positions are indicated. Orientations of N-termini (N^term^) and C-termini (C^term^) are shown in the corners. **(b-e)** EnFOE-induced accumulation of reactive oxygen species (ROS) measured in leaf discs from *Nicotiana benthamiana* transiently expressing *Arabidopsis thaliana* MIK2 (*At*MIK2.1) or *Solanum lycopersicum* MIK2-A (*Sl*MIK2-A). Proteins were expressed as wildtype or with indicated amino acids substitutions. Recorded values were normalized to background and mock-treated samples and summed up over a period of 60 min (cumulative [cum.] ROS) and are given in relative light units (rlu). To enhance comparability, all individual values were standardized to average values of wildtype controls within each independent experiment (collective values of wildtype average at 1.0). Data are pooled from 3-5 independent experiments, each with n = 4-8, bars indicate mean and standard deviation. Asterisks indicate statistically significant differences compared to respective wildtype with **** = *P* < 0.0001, *** = *P* < 0.001, ** = *P* < 0.01, * = *P* < 0.05, ns = not significant with *P* > 0.05, ordinary one-way ANOVA with Dunnett’s multiple comparisons test. **(f)** Cumulative cytosolic elevation of calcium ion concentration (cum. [Ca^2+^]cyt) in *Arabidopsis thaliana* Col-0^AEQ^ wildtype and *mik2*^AEQ^ mutant seedlings after elicitation with EnFOE [0.5 µg/mL] unbound. Unbound fractions resulted from filtering of EnFOE preparations through columns lined with the ectodomains of *Arabidopsis thaliana* MIK2 (e*At*MIK2) wildtype, e*At*MIK2 with indicated alanine-substitutions or empty column controls that were only loaded with apoplastic wash fluids (AWFs). Data shown are pooled from three independent experiments, each with n=4. Values of individual seedlings (each plotted dot) were summed over a period of 30 min after elicitation. Relative calcium levels are presented as luminescence counts per second relative to total luminescence counts remaining (L/Lmax). Bars indicate mean and standard deviation. Letters indicate significant differences (one-way ANOVA, Tukey’s test, *P* < 0.001). **(g)** Western blot visualization of C-terminally HA-tagged proteins in eluted fractions corresponding to data of one experiment from fig.6f. Blots were probed with α-HA antibodies. Approximate protein size is marked on the left edge and given in kilodaltons (kDa).

To test whether the identified conserved residues might be involved in the interaction between the *Fusarium* elicitor and MIK2, we compared the ligand-depleting activity of wildtype and mutant ectodomains using our ligand-depletion assay. As previously described, e*At*MIK2^wildtype^ fully depleted elicitor-activity from EnFOE preparations, with unbound fractions not inducing a significant MIK2-dependent calcium transient in Col-0^AEQ^ wildtype seedlings (**fig. 6f-g**). In contrast, the ectodomain of the tested triple substitution mutant e*At*MIK2^Y318A/D367A/Q369A^ completely lost the ability to deplete elicitor-activity from EnFOE, with unbound fractions inducing a calcium response that does not significantly deviate from that of fractions run through control columns that did not contain any ectodomains (**fig. 6f-g; fig.S7m**). We also tested the corresponding triple and a quadruple mutant of e*Sl*MIK2-A. However, in contrast to the respective full-length proteins (**fig.S5f, S5i; fig.S7f, S7i**), the expression and purification of the mutant ectodomains were repeatedly weaker than that of the wildtype protein, making a quantitative comparison between the ligand-depleting activity impossible (**fig.S5m**).

Nevertheless, these results suggest a potentially common EnFOE-binding interface on *At*MIK2 and *Sl*MIK2-A, which, in *Arabidopsis*, might be extended N-terminally towards the SCOOP binding-site on *At*MIK2 or dependent on its structural features.

## Discussion

### *Fusarium* elicitor-responsiveness and MIK2-clade proteins are conserved across plant families

The *A. thaliana At*MIK2 receptor integrates endogenous and exogenous peptide signaling. Most *Atmik2* mutant phenotypes, including reduced salt tolerance (Julkowska *et al*., 2016; van der Does *et al*., 2017), impaired cell wall damage responses (van der Does *et al*., 2017), and increased herbivore susceptibility (Stahl *et al*., 2022), link directly or indirectly to its role as receptor for SCOOP peptides (Yang *et al*., 2023a; Zhai *et al*., 2024) . In contrast, enhanced susceptibility to *Fusarium* (van der Does *et al*., 2017; Coleman *et al*., 2021) cannot be attributed to SCOOP-MIK2 signaling (Yang *et al*., 2023a), supporting a SCOOP-independent function of *At*MIK2 in sensing *Fusarium*-derived signals.

Motifs with similarity to plant SCOOPs occur in various *Fusarium* proteins, and synthetic SCOOP-LIKE (SCOOPL) peptides induce PTI responses in *Arabidopsis* (Hou *et al*., 2021; Rhodes *et al*., 2021) . However, SCOOP peptides are exclusive to the Brassicales, and SCOOP-responsiveness is restricted accordingly (Gully *et al*., 2019; Snoeck *et al*., 2024) . Similarly, non-Brassicales species fail to perceive SCOOPLs (**fig. 1b-d**). In contrast, EnFOE elicits responses across distant species including soybean, tomato, and barley (Coleman *et al*., 2021) (**fig. 1a**). This may reflect additional elicitor-active molecules in EnFOE or might show that SCOOPLs are not the primary source of MIK2-dependent elicitor-activity. Even endogenous SCOOPs remain largely based on predictions and might differ substantially from their native bioactive forms (Guillou *et al*., 2022) .

Peptide sequence and structure profoundly affect interspecies recognition. For example, the flg22 peptide epitope is perceived by *Arabidopsis* and tomato, whereas flg15 is recognized only by tomato (Felix *et al*., 1999) . Similarly, small cysteine-rich protein (SCP) perception depends on tertiary structure in *Arabidopsis,* but linear fragments are sufficient for other members of the *Brassicaceae* (Yang *et al*., 2023b) . Thus, identifying the active EnFOE elicitor requires biochemical validation beyond bioinformatics.

Although SCOOP perception is Brassicales-specific (Snoeck *et al*., 2024), MIK2-clade genes are conserved across seed plants with lineage-specific patterns of duplication and retention (**fig. 1e-f; fig.S2c**). Such dynamics likely reflect adaptations to distinct environmental conditions and pathogen pressures. Neofunctionalization of duplicated receptors may be advantageous under high pathogen pressure, whereas receptors may be lost when constraints relax. Such dynamics have been described for the INR receptor in legumes (Snoeck *et al*., 2022) . Interestingly, the Brassicales retain fewer MIK2-clade genes than species of other plant orders (**fig. 1e-f; fig.S2c**), potentially reflecting constraints imposed by SCOOP-mediated signaling, where additional copies could disrupt finely balanced signaling outputs.

Notably, *N. benthamiana* is unresponsive to EnFOE. Whether this reflects the absence of a functional MIK2-clade receptor remains unresolved, as its genome is incompletely annotated. Even though two *N. benthamiana* proteins might share a common origin with *At*MIK2 according to our phylogenetic analysis, they form a divergent cluster away from other Solanales proteins. They also do not return to *At*MIK2 or tomato MIK2-clade proteins as top hits when reciprocally sequence-compared against respective proteomes. Taken together, the broad occurrence of EnFOE responsiveness and MIK2-clade proteins suggests a shared *Fusarium* elicitor perception mechanism across plant orders.

### A MIK2-clade protein from *Solanum lycopersicum* shows similar properties to *At*MIK2 in *Fusarium* elicitor perception

Because MIK2-clade genes exhibit strong species-specific duplication and retention patterns (**fig. 1e-f; fig.S2c**), identifying a putative *At*MIK2 orthologue based on phylogeny was not straightforward. We therefore tested all five MIK2-clade proteins and the MIK2-sister-clade protein from tomato. Ectopic expression of *Sl*MIK2-A conferred EnFOE-sensitivity to *Nicotiana benthamiana* comparable to *At*MIK2 (**fig. 2c**). Consistent with the lack of responsiveness of tomato to *At*SCOOP12 and *Foc*SCOOPL peptides (**fig. 1b-d**), none of the tomato proteins conferred sensitivity to respective peptides (**fig. 2b; fig.S3b-c**). This suggests similar mechanisms for EnFOE perception involving MIK2-clade proteins across plant orders, while SCOOP perception remains Brassicales-specific (Snoeck *et al*., 2024) . Further evidence that *At*MIK2 and *Sl*MIK2-A derive from a common ancestor is provided by a synteny analysis (**fig. 2d**). Both genes reside at a conserved genomic region in respective plant species. Notably, the syntenic locus in tomato additionally harbors genes coding for *Sl*MIK2-B, -C and -D (**fig. 2d**).

*SlMIK2-A* generates four splice-variants differing in ecto- or kinase domains (**fig. 2e**). Splice-variant #1 is predominantly expressed (**fig. 2e; fig.S3e**) and the only one translated into a protein that mediates the perception of EnFOE (**fig. 2f**). The observed alternative splicing might play a role in the fine-tuning of plant immune responses, as previously described (Sanabria & Dubery, 2016; Kufel *et al*., 2022; Godinho *et al*., 2024) . As often observed for the expression of PRRs when plants are exposed to MAMPs or pathogen challenge (Zipfel *et al*., 2004; Hou *et al*., 2021), *SlMIK2-A* (splice-variant #1) expression is induced by EnFOE or chitin treatment and by *Fusarium* infection (**fig. 2h**).

### *Sl*MIK2-A forms EnFOE-induced complexes with tomato BAK1 orthologues and depletes elicitor-activity from EnFOE

MAMP-perception by LRR-type PRRs often involves ligand-induced complex formation with SERK family co-receptors (Chinchilla *et al*., 2007; Heese *et al*., 2007; Roux *et al*., 2011; Peng & Kaloshian, 2014; Wei *et al*., 2020) . Likewise, SCOOP and EnFOE promote association between *At*MIK2 and *At*BAK1 (Coleman *et al*., 2021; Hou *et al*., 2021; Rhodes *et al*., 2021) . Similarly, *Sl*MIK2-A forms EnFOE-induced complexes with tomato BAK1 orthologues, while *At*SCOOP12 or *Foc*SCOOPL^FOXB_11846^do not promote interactions between the tomato pairs (**fig. 3a-b; fig.S4a**). This provides indirect evidence for an interaction between *Sl*MIK2-A and a component of EnFOE and suggests a similar mechanism involving ligand-induced complex formation with SERK-family proteins for EnFOE-perception across plant orders.

Additional evidence for an interaction between an elicitor-active component of EnFOE and *At*MIK2 and *Sl*MIK2-A is provided by a modified ligand-depletion assay (Shu *et al*., 2026) . The ectodomains of *At*MIK2 and *Sl*MIK2-A nearly completely depleted elicitor-activity from EnFOE (**fig. 3e**). The fact that this was also genetically *At*MIK2-dependent when *Arabidopsis* was challenged by the tomato *Sl*MIK2-A-depleted elicitor fraction, indicates that both ectodomains might interact with similar elicitor molecules. In contrast, *At*SCOOP12 is only depleted by the ectodomain of *At*MIK2 **(fig. 3d**). The specificity of this assay is further supported because flg22 preparations retained full *At*FLS2-dependent activity after passage through e*At*MIK2-, e*Sl*MIK2-A- and e*Sl*MIK2-B-lined columns, while being depleted by the ectodomain of the cognate receptor *At*FLS2 (**fig. 3f**). These results additionally validate the adapted depletion assay as a tool to probe receptor-ligand interactions. Unlike many *in vitro* approaches (Sandoval & Santiago, 2020), this assay is simple and suited for studying interactions when ligands are unknown or not available in purified form, such as EnFOE. Nevertheless, the assay has its limitations. It depends on successful *in planta* expression of intact ectodomains in sufficient amounts and a sensitive readout. It may fail to detect weak or multicomponent interactions and therefore requires case-to-case optimization and appropriate controls. Additionally, when based on a readout that depends on *Arabidopsis*, as we have it here, there is a chance of missing responses to elicitors that this plant species does not perceive. Given the observation that *Slmik2-A*^CRISPR^ mutants are still partially responsive to treatment with EnFOE (**fig. 4a, 4c**), suggest that the elicitor fraction contains elicitor-active compounds, which are differentially perceived by *Arabidopsis* and tomato. Hence, a potential interaction between e*Sl*MIK2-A and a divergent or completely different elicitor-active molecule might go unnoticed.

### *Sl*MIK2-A is important for EnFOE-induced PTI responses and resistance to *Fusarium oxysporum* infection in tomato

Knockout of *SlMIK2- A* clearly reduced PTI responses to EnFOE. However, unlike the complete insensitivity of *Atmik2* mutants (Coleman *et al*., 2021), *Slmik2- A* mutants retained residual responses to EnFOE (**fig. 4a, 4c**). This may reflect that the EnFOE enrichment protocol was optimized for *Arabidopsis* to remove non-*At*MIK2-dependent activity, whereas tomato might perceive an additional elicitor not recognized by *Arabidopsis*, potentially *via* a PRR that is not conserved across plant lineages (Zipfel *et al*., 2006; Ranf *et al*., 2015; Wang *et al*., 2016) . Alternatively, a different epitope of the same elicitor could be sensed by a distinct receptor, as seen for *Sl*FLS2 and *Sl*FLS3 (Hind *et al*., 2016) . Future efforts to identify the specific EnFOE molecule triggering MIK2-dependent PTI responses will clarify this. Although no other tomato MIK2-clade protein conferred EnFOE-sensitivity to *N. benthamiana* (**fig. 2c**), a role in the native tomato system remains possible.

Surprisingly, *Slmik2- A* mutants showed no difference in chitin-triggered ROS production compared to wildtype (**fig. 4b**), but chitin-induced ethylene accumulation was reduced (**fig. 4d**). However, flg22-induced ethylene remained unaffected, indicating that the mutants are not broadly impaired in MAMP-triggered ethylene responses (**fig. 4e**). Similarly, loss of *At*MIK2 reduces responses to certain MAMPs (Rhodes *et al*., 2021) and *At*MIK2 may act as a scaffolding component coordinating perception events in *Arabidopsis* (Delplace *et al*., 2024) . Loss of *Sl*MIK2-A may have comparable effects on certain PTI-pathways in tomato. Overall, these results demonstrate an important contribution of *Sl*MIK2-A to EnFOE-induced PTI responses in tomato.

The reduced EnFOE-responsiveness of *Slmik2-A*^CRISPR^ mutants is accompanied by enhanced susceptibility to infection (**fig. 4f-g**). We used the *F. oxysporum* f.sp. *lycopersici* strain *Fol007*, which only mildly affects wildtype plants. *Fol* strains can suppress PTI signaling *via* effector proteins, such as Avr2, which inhibits EnFOE-triggered responses in *Arabidopsis* (Di *et al*., 2017; Tintor *et al*., 2020; Coleman *et al*., 2021; Lamo *et al*., 2021; Blekemolen *et al*., 2023) . Consequently, the contribution of a single RK to resistance can be small if EnFOE-induced PTI is already partially suppressed by the pathogen. Even without pathogen interference, loss of one PRR can have a limited effect due to redundant MAMP perception (Zipfel *et al*., 2004; Boller & Felix, 2009; Fan *et al*., 2022) . Contributions of individual PRRs to overall resistance can also be diminished when the pathogens are supported to overcome the first layers of pre-invasive resistance (Zipfel *et al*., 2004) . Despite facilitating *Fol007* infection by damaging of roots, *Slmik2-A*^CRISPR^ mutants showed significantly greater reductions in plant height and fresh-weight than wildtype plants (**fig. 4f-g**), highlighting the contribution of *Sl*MIK2-A to quantitative disease resistance in tomato.

### Conserved residues on the ectodomains of *At*MIK2 and *Sl*MIK2-A are important for conferring EnFOE-sensitivity

Our mutational analysis showed that residues essential for SCOOP-binding in *At*MIK2 (Jia *et al*., 2024; Snoeck *et al*., 2024; Wu *et al*., 2024) are also required for EnFOE-induced responses, suggesting that *Fusarium* elicitor perception could also be mediated by this interface or may require its structural integrity (**fig. 5b-c**). Notably, the structural equivalents of the relevant residues in *At*MIK2 are not conserved in *Sl*MIK2-A, and substitution mutations in the tomato receptor did not impair EnFOE perception (**fig. 5a, 5d**). This indicates an at least partially distinct interaction surface, despite overall functional similarities between the receptors. Our data further highlight a central role of *At*MIK2 glycosylation in EnFOE perception. The *N*-glycosylation site *At*MIK2^N410^, important for *At*SCOOP12-induced interaction with the co-receptor *At*BAK1, is also essential for conferring EnFOE-sensitivity to *N. benthamiana* and for EnFOE-induced receptor complex formation (**fig. 5e-f**). The absence of a conserved N410 equivalent in *Sl*MIK2-A and the lack of significant effect of alternative glycosylation-site mutations suggest that different structural solutions could support receptor function and co-receptor engagement in those two species (**fig. 5g-h**). *Sl*MIK2-A might have evolved alternative structural arrangements that bypass the glycan-dependent activation mechanism required in *Arabidopsis*. While *N*-glycosylation has been commonly reported to be important for receptor maturation, plasma membrane targeting and accumulation (Nekrasov *et al*., 2009; Saijo *et al*., 2009; Häweker *et al*., 2010; Chen *et al*., 2020), a direct involvement in ligand-receptor-coreceptor interaction has so far only been reported for SCOOP/SCOOPL-MIK2-BAK1 complexes (Jia *et al*., 2024; Wu *et al*., 2024), potentially representing a Brassicales-specific innovation in MIK2-clade receptors.

By focusing on residues conserved between *At*MIK2 and *Sl*MIK2-A but absent in the closely related non-functional paralogue *Sl*MIK2-B, we identified a cluster of amino acids (Y318/310, D367/359, Q369/361, and, to a lesser extent, K345/337) that are specifically required for EnFOE-induced responses in both species (**fig. 6a-e; fig.S6**). Notably, mutation of these residues had little to no effect on SCOOP responsiveness in single mutants, indicating that they define a functionally distinct surface (**fig.S6**). The strong impact of combinatorial mutations suggests cooperative contributions to ligand perception or receptor activation (**fig. 6c; fig.S6**). The partial effect of Y318 on SCOOP-induced signaling in one of the double mutants and all triple mutants that contained the Y318A substitution is consistent with structural data showing interaction of the C-terminal region of *At*MIK2 with the side chain of this residue (Jia *et al*., 2024; Wu *et al*., 2024), implying that this site may represent a point of convergence between different ligand recognition modes.

The ligand-depletion assay provides further indications for a role of these residues in binding the elicitor-activity in EnFOE. The inability of the corresponding mutated ectodomain to deplete elicitor-activity strongly suggests that this surface contributes to ligand interaction rather than merely downstream signaling (**fig. 6f**). Although technical limitations prevented a similar quantitative analysis for *Sl*MIK2-A ectodomain mutants, the consistency between full-length receptor phenotypes and *At*MIK2 ectodomain behavior supports the existence of a conserved functional interface.

Taken together, our findings support a new hypothesis in which *At*MIK2 and *Sl*MIK2-A perceive EnFOE through a shared but evolutionarily flexible surface that is spatially linked to, yet partially independent of the SCOOP-binding interface. In *Arabidopsis*, this interface may extend towards or structurally depend on the SCOOP-binding region, whereas in tomato, alternative residues could compensate for the lack of conservation. These differences may reflect species and receptor-specific adaptation to ligand diversity, similar to what has been reported for flagellin-derived immunogenic patterns (Trdá *et al*., 2014; Cheng *et al*., 2021; Colaianni *et al*., 2021) . Additionally, the evolution of MIK2 as a SCOOP receptor in the Brassicales likely enforced different constraints on the binding site for the *Fusarium* elicitor when compared with the tomato receptor, which did not have to accommodate or sustain this additional function.

We here present insights into the evolutionary trajectory of a multifunctional plant PRR. *Arabidopsis At*MIK2 perceives the exceptionally diverse family of endogenous SCOOP peptides while simultaneously being involved in the recognition of a yet-to-be-identified MAMP from *Fusarium* and related fungi (Coleman *et al*., 2021; Hou *et al*., 2021; Rhodes *et al*., 2021; Snoeck *et al*., 2024) . There are a few examples of cell-surface receptors with dual ligands, including *Arabidopsis* CARD1/HPCA1 (Laohavisit *et al*., 2020; Wu *et al*., 2020), potato NILR1 (Huang *et al*., 2023; Huang *et al*., 2024) or *Arabidopsis* RLP30 (Yang *et al*., 2023b) . Similar flexibility can also be observed for certain intracellular receptors that can perceive different microbial effector molecules (Brabham *et al*., 2024; Kunz *et al*., 2025) . On the other hand, there are examples of specific ligands that are perceived by multiple redundant or distantly related receptors in *Arabidopsis*, as is the case for IDA, which is perceived by the receptors HAE and HSL2 (Butenko *et al*., 2003; Cho *et al*., 2008; Stenvik *et al*., 2008; Butenko *et al*., 2014) ; the CEP4 peptide, which is perceived by CEPR1, CEPR2 and RLK7 in a tissue-specific manner (Rzemieniewski *et al*., 2024) ; PEP1 and PEP2, which are both perceived by PEPR1 and PEPR2 (Huffaker *et al*., 2006; Yamaguchi *et al*., 2010) ; as well as PEP7, which has been shown to interact with the ectodomain of PEPR1 *in vitro* but has also been identified as ligand for the unrelated receptor kinase SIRK1 (Tang *et al*., 2015; Wang *et al*., 2022) . It remains an interesting question why *At*MIK2 evolved the exclusive ability to function in the perception of SCOOPs and the *Fusarium* elicitor, and why potential gene duplications that could facilitate the distribution of subfunctions were not retained in most members of the Brassicales. The fact that SCOOPs and their perception are restricted to members of the Brassicales, while genes of the MIK2-clade and *Fusarium* elicitor-responsiveness are commonly found in angiosperms (**fig. 1a-f**) could suggest a more ancient role in the perception of the microbial signature, while the function as receptor for the endogenous peptides has been gained later, either simultaneously or after the appearance of SCOOPs themselves. Situations where the perception of phytocytokines might have evolved through the molecular mimicry of microbial signatures have been proposed before (Hou *et al*., 2021; Ngou *et al*., 2024) . Intriguingly, there is accumulating evidence that proteins of the MIK2-clade outside the order Brassicales play a direct or indirect role in the perception of MAMPs or in pathogen resistance (Bañales *et al*., 2025; Zhu *et al*., 2025) . Finally, it is also conceivable that SCOOPs initially evolved to serve different functions, for example as antimicrobial peptides. Such activities have been demonstrated for multiple SCOOPs, even before they were designated as such (Yadeta *et al*., 2014; Neukermans *et al*., 2015; Yu *et al*., 2020; Zhang *et al*., 2022) . It is tempting to speculate that one or multiple Brassicales-specific mutations on the ectodomain of MIK2 might then have given rise to the interaction with SCOOPs, while still retaining the initial function in perception of the *Fusarium* elicitor.

## Acknowledgements

This work was supported by the grant ”Fusarium elicitor-triggered immunity in model and crop plants” from the German Research Foundation (project number: 547682052) (R.H.), the Gatsby Charitable Foundation (C.Z.), the European Research Council under the European Union’s Horizon 2020 research and innovation program no. 773153 (project ‘IMMUNO-PEPTALK’) (C.Z.), the Swiss National Science Foundation grant no. 320030_228294 (C.Z.), the University of Zurich (C.Z. and S.S.), the Uniscientia Foundation (C.Z.), the de.NBI Cloud within the German Network for Bioinformatics Infrastructure (de.NBI) and ELIXIR-DE (Forschungszentrum Jülich and W-de.NBI-001, W-de.NBI-004, W-de.NBI-008, W-de.NBI-010, W-de.NBI-013, W-de.NBI-014, W-de.NBI-016, W-de.NBI-022) (C.R.), and the German Research Foundation in frame of the collaborative research center SFB924 (C.S. and Y.R.).

We thank Frank L. W. Takken for providing the *Fusarium oxysporum* f.sp. *lycopersici* strain *Fol*007 and Stefanie Ranf-Zipproth for providing the GoldenGate-based cloning system. In addition, we especially thank Jonas Schweigel, Julia Hagn, Lena Glanz, Samuel Sadorf, Kristina Freymark, and Jakob Dietz, who set the basis for certain experiments with preliminary results from their student theses.

## Author contributions

J.M. and R.H. designed the study. J.M. performed experiments, analysed and visualised data. Y.R. generated tomato CRISPR mutants. C.R. analysed and visualised the phylogeny and representation of gene copy numbers. S.S. analysed and visualised the synteny. C.S., C.Z. and R.H. supervised the study and acquired funding. J.M., R.H., C.R., and S.S. wrote the manuscript. All authors reviewed, edited and approved the final manuscript.

## Data availability

All data is deposited in a public repository and can be found under the following link: https://doi.org/10.5281/zenodo.20807889

## Supporting Information

The following Supporting Information is available for this article:

**fig.S1** Visualization of predicted local distance difference test (pLDDT) scores for AF3 predictions.

**fig.S2** Elicitor-responsiveness and phylogeny of *MIK2-CLADE* genes in plants of different orders.

**fig.S3** Analysis of MIK2-clade proteins from *Solanum lycopersicum*.

**fig.S4** Ligand-induced receptor-complex formation, expression of affinity-tagged ectodomains and schematics of *Atmik2*^AEQ^ and *Slmik2-A* CRISPR mutants.

**fig.S5** Western blot visualization of C-terminally cMyc-tagged proteins.

**fig.S6** Influence of conserved residues on EnFOE-sensitivity conferred by *At*MIK2 and *Sl*MIK2-A.

**fig.S7** Quantification of Western blot signals of C-terminally cMyc-tagged proteins.

**table S1a** list of cloned constructs

**table S1b** primer list

**table S1c** guideRNAs for CRISPR/Cas9-mediated mutagenesis

**table S1d** plant species for phylogeny

**table S1e** phylogeny sequences

**table S1f** phylogeny copy numbers

