## Supplementary Figures for "A tomato MIK2-clade receptor is involved in the perception of a *Fusarium*-derived elicitor"

The following Supporting Information is available for this article:

**fig.S1** Visualization of predicted local distance difference test (pLDDT) scores for AF3 predictions.

**fig.S2** Elicitor-responsiveness and phylogeny of *MIK2-CLADE* genes in plants of different orders.

**fig.S3** Analysis of MIK2-clade proteins from *Solanum lycopersicum*.

**fig.S4** Ligand-induced receptor-complex formation, expression of affinity-tagged ectodomains and schematics of *Atmik2*$^{AEQ}$ and *Slmik2-A* CRISPR mutants.

**fig.S5** Western blot visualization of C-terminally cMyc-tagged proteins.

**fig.S6** Influence of conserved residues on EnFOE-sensitivity conferred by *At*MIK2 and *Sl*MIK2-A.

**fig.S7** Quantification of Western blot signals of C-terminally cMyc-tagged proteins.

**table S1a** list of cloned constructs

**table S1b** primer list

**table S1c** guideRNAs for CRISPR/Cas9-mediated mutagenesis

**table S1d** plant species for phylogeny

**table S1e** phylogeny sequences

**table S1f** phylogeny copy numbers

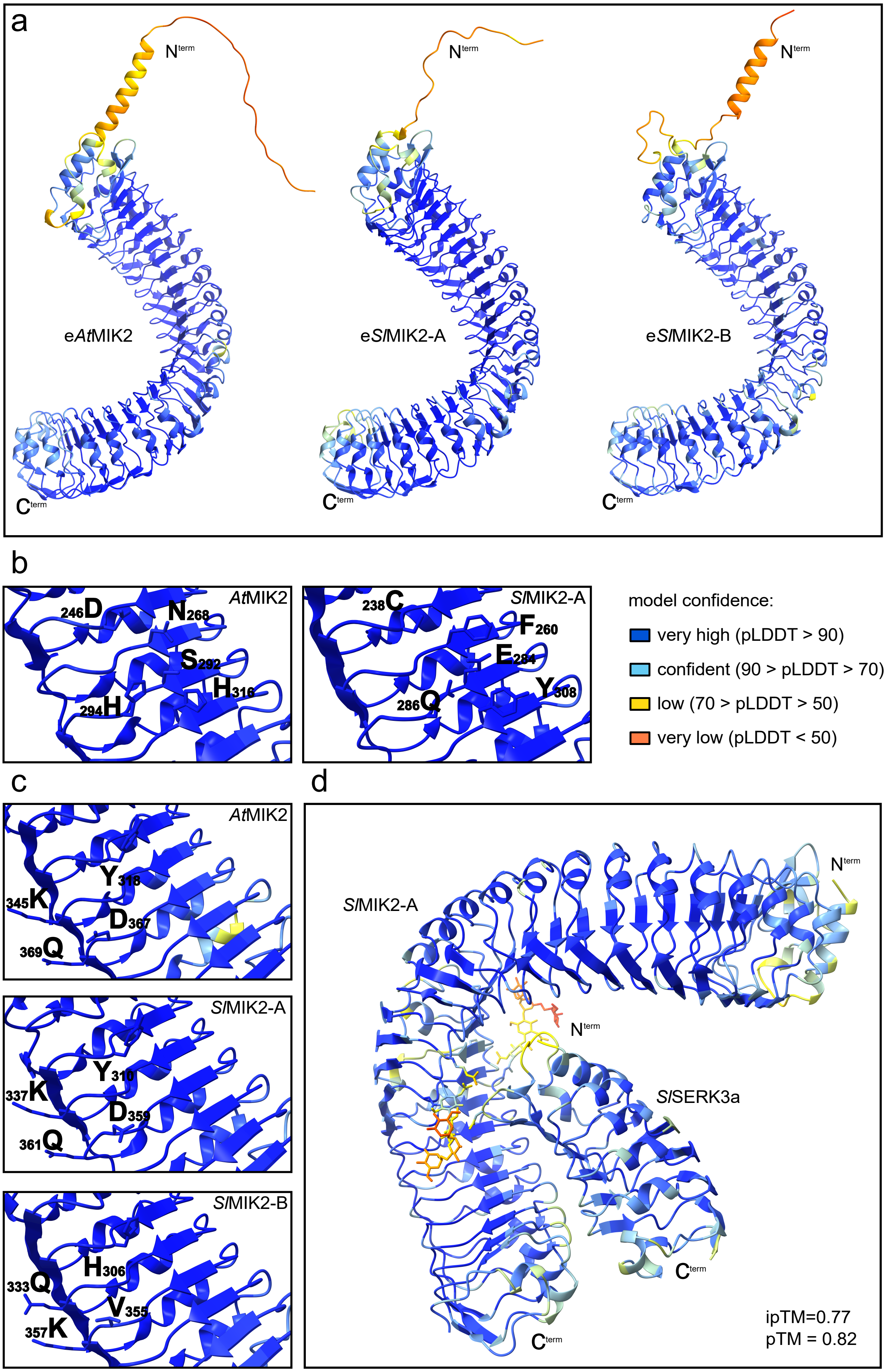
a
Nterm
eAtMIK2
eSlMIK2-A
eSlMIK2-B
Cterm
b
AtMIK2
246D
N268
S292
H316
294H
SlMIK2-A
238C
F260
E284
Y308
286Q
model confidence:
very high (pLDDT > 90)
confident (90 > pLDDT > 70)
low (70 > pLDDT > 50)
very low (pLDDT < 50)
c
AtMIK2
Y318
345K
D367
369Q
SlMIK2-A
Y310
337K
D359
361Q
SlMIK2-B
H306
333Q
V355
357K
d
SlMIK2-A
SlSERK3a
ipTM=0.77
pTM = 0.82

**fig.S1 | Visualization of predicted local distance difference test (pLDDT) scores for AF3 predictions.** AlphaFold produces a per-residue confidence score (pLDDT) between 0 and 100, with higher values indicating higher confidence. Colors represent pLDDT values according to the legend. **(a)** pLDDT visualization of ectodomains of *At*MIK2 (Uniprot ID: Q8VZG8), *Sl*MIK2-A (Uniprot ID: A0A3Q7G8D9) and *Sl*MIK2-B (model in supplementary material). Orientation of N- and C-termini are indicated ($N^{term}$, $C^{term}$). **(b)** Close-up of pLDDT visualization of *At*MIK2 (left) and *Sl*MIK2-A (right). Important SCOOP-binding residues (Jia *et al.,* 2024; Snoeck *et al.,* 2024; Wu *et al.,* 2024) on *At*MIK2 and their structural equivalents (same amino acid positions according to structural alignment) on *Sl*MIK2-A are highlighted and respective amino acids and positions are indicated. **(c)** Close-up of pLDDT visualization of *At*MIK2 (top), *Sl*MIK2-A (middle), and *Sl*MIK2-B (bottom). Aligned surface-exposed residues conserved between *At*MIK2 and *Sl*MIK2-A, but not *Sl*MIK2-B, are highlighted and respective amino acids and positions are indicated. **(d)** pLDDT visualization of AlphaFold-Multimer (AFM)-predicted complex structure of *Solanum lycopersicum* MIK2-A (*Sl*MIK2-A) and SERK3a (*Sl*SERK3a) ectodomains including glycan chains attached to *Sl*MIK2-$A^{N431}$ and *Sl*MIK2-$A^{N471}$. Orientation of N-termini ($N^{term}$) and C-termini ($C^{term}$) are shown in the corners. The predicted template modeling (pTM) score and the interface predicted template modeling (ipTM) score are shown on the right bottom.

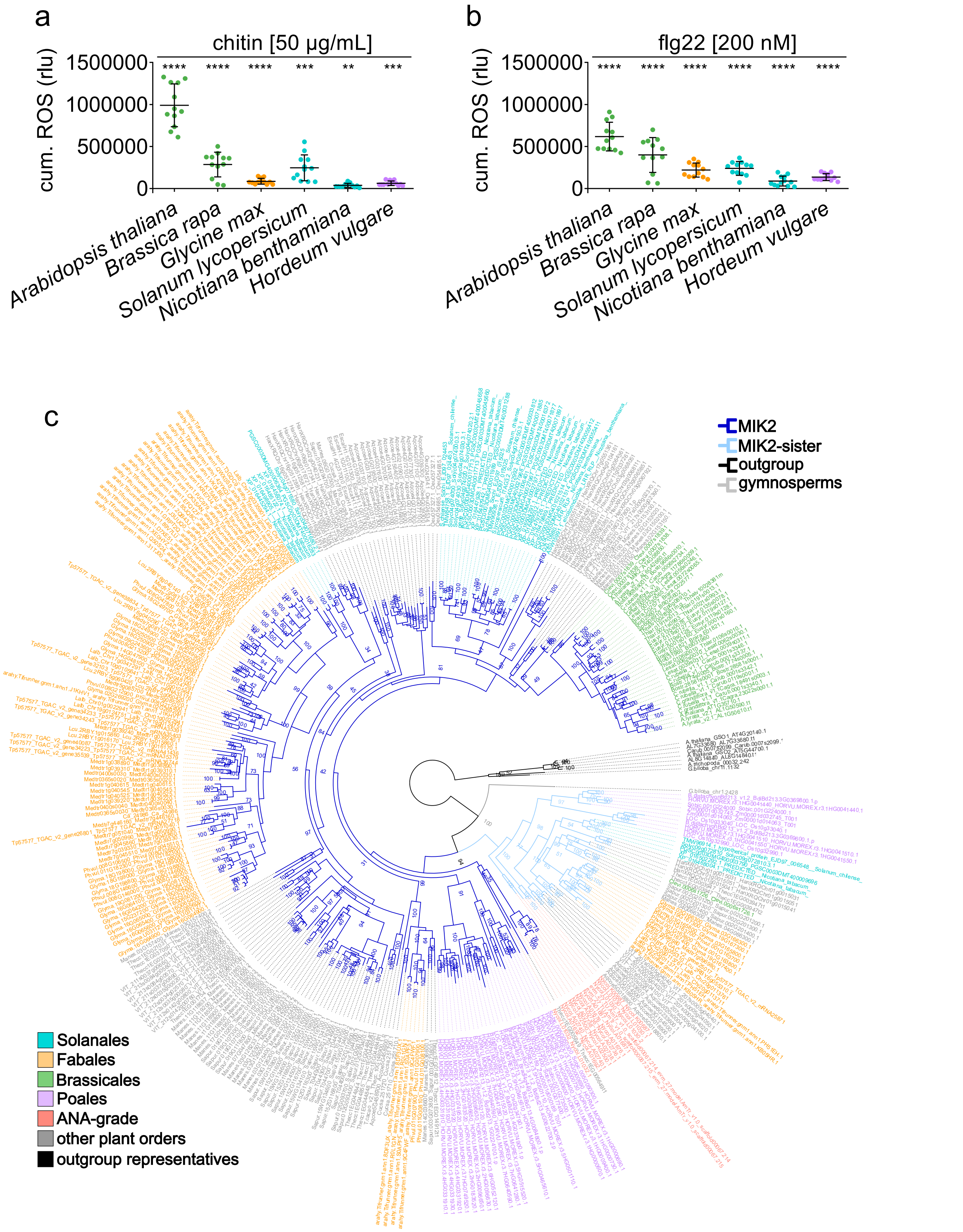


**fig.S2 | Elicitor-responsiveness and phylogeny of *MIK2-CLADE* genes in plants of different orders. (a-b)** Accumulation of reactive oxygen species (ROS) measured in leaf discs from Brassicales species *Arabidopsis thaliana* and *Brassica rapa* (Pak choi), Fabales species *Glycine max* (soybean), Solanales species *Solanum lycopersicum* (tomato) and *Nicotiana benthamiana,* and Poales species *Hordeum vulgare* (barley) following elicitation with 50 µg/mL chitin or 200 nM flg22. Recorded values were normalized to background and mock-treated samples and summed over a period of 60 min (cumulative [cum.]

ROS) and are given in relative light units (rlu). Data are pooled from 3 independent experiments, each with n = 4, bars indicate mean and standard deviation. Asterisks indicate statistically significant differences from respective mock-treated samples. Unpaired t-test was performed on data before normalization to mock-treated samples. **** = $P < 0.0001$, *** = $P < 0.001$, ** = $P < 0.01$. **(c)** Maximum likelihood (ML) phylogeny of *MIK2* genes including representatives of major angiosperm lineages and Ginkgo (gymnosperm). An early duplication event separated the MIK2-clade (dark blue branches) from the MIK2-sister-clade (light blue branches). Genes are color-coded according to plant orders Fabales (yellow), Brassicales (green), Solanales (turquoise), and Poales (lilac). ANA-grade (red) refers to basal angiosperm orders including Amborellales, Nymphaeales, and Austrobaileyales. Genes of species from other plant orders are colored in gray, outgroup representatives are colored in black. Numbers indicate bootstrap support in %.

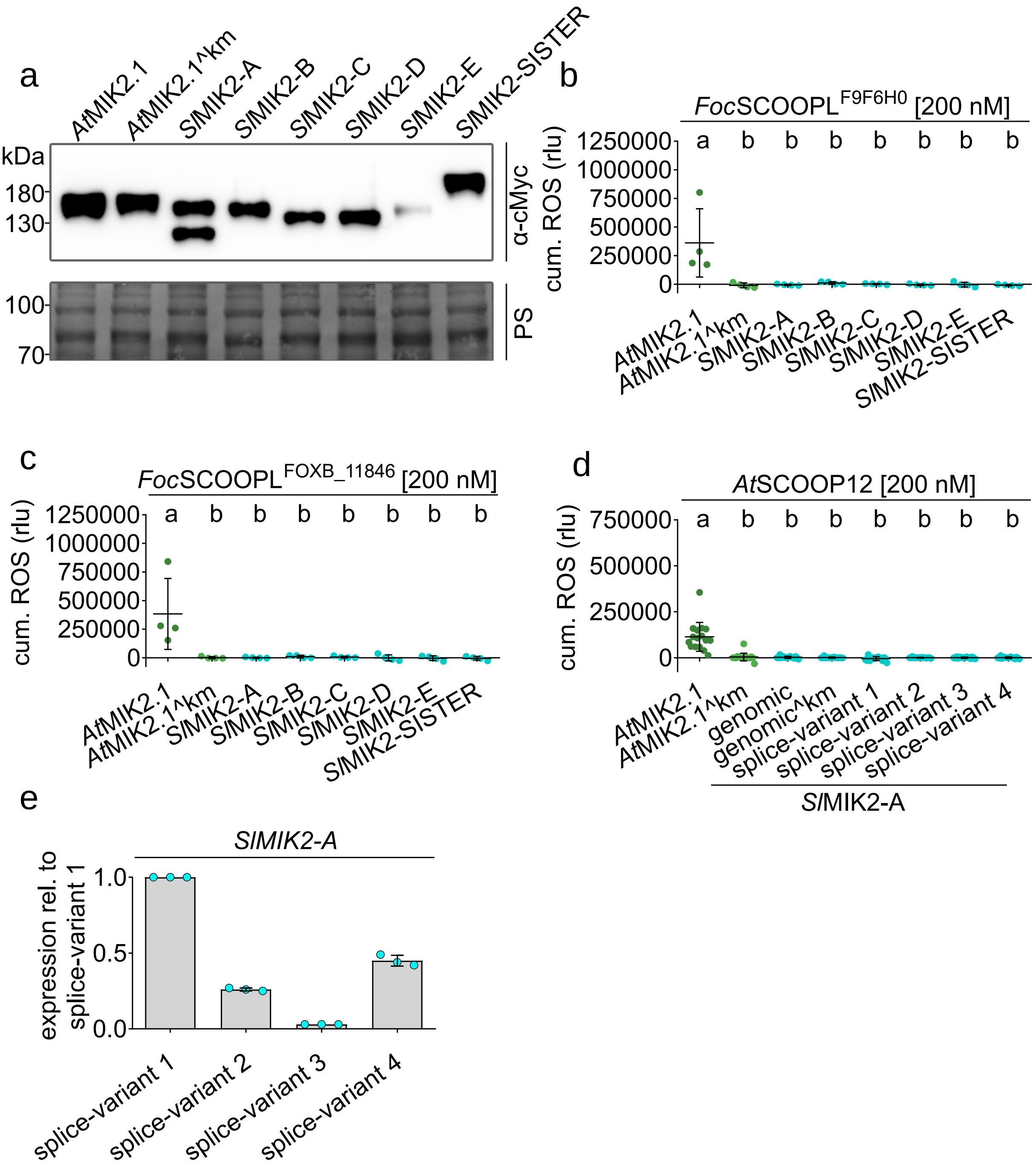


**fig.S3 | Analysis of MIK2-clade proteins from *Solanum lycopersicum*. (a)** Western blot visualization of C-terminally cMyc-tagged proteins corresponding to data of one experiment from fig.2b-c. Blots were probed with α-cMyc antibodies. Approximate protein size is marked on the left edge and given in kilodaltons (kDa). Equal protein loading was confirmed by Ponceau S staining (PS). **(b-c)** *Foc*SCOOPL$^{F9F6F0}$- and *Foc*SOOPL$^{FGSG_07177}$-induced accumulation of reactive oxygen species (ROS) measured in leaf discs from *Nicotiana benthamiana* transiently overexpressing *Arabidopsis thaliana* MIK2.1 (*At*MIK2), a kinase-inactive version of *At*MIK2.1 (^km, K802A) or *Solanum lycopersicum* proteins from the MIK2-clade and MIK2-sister-clade (*Sl*MIK2-A to E, *Sl*MIK2-SISTER). Recorded values were normalized to background and mock-treated samples and summed over a period of 60 min (cumulative [cum.] ROS) and are given in relative light units (rlu). Data show one independent experiments, n = 4. Bars indicate mean and standard deviation. Letters show significant differences (one-way ANOVA, Tukey's test, $P < 0.001$). **(d)** *At*SCOOP12-induced accumulation of reactive oxygen species (ROS) measured in leaf discs from *Nicotiana benthamiana* transiently overexpressing *At*MIK2.1, *At*MIK2.1^km and different versions of *Sl*MIK2-A (genomic sequence, K791A mutated ^km version and coding sequence of individual splice variants). Recorded values were normalized to background and mock-treated samples and summed over a period of 60 min (cum. ROS) and are given in relative light units (rlu). Data are pooled from 4 independent experiments, each with n = 4, bars indicate mean and standard deviation. Letters show significant differences (one-way ANOVA, Tukey's test, $P < 0.0001$). **(e)** Expression of individual *SlMIK2-A* splice-variant transcripts in *Solanum lycopersicum* cv. M82 wildtype plants. Values were normalized to the expression of the housekeeping gene *SlEF1α* and are shown as expression relative to splice-variant 1. Individual dots show results from 3 biological replicates.

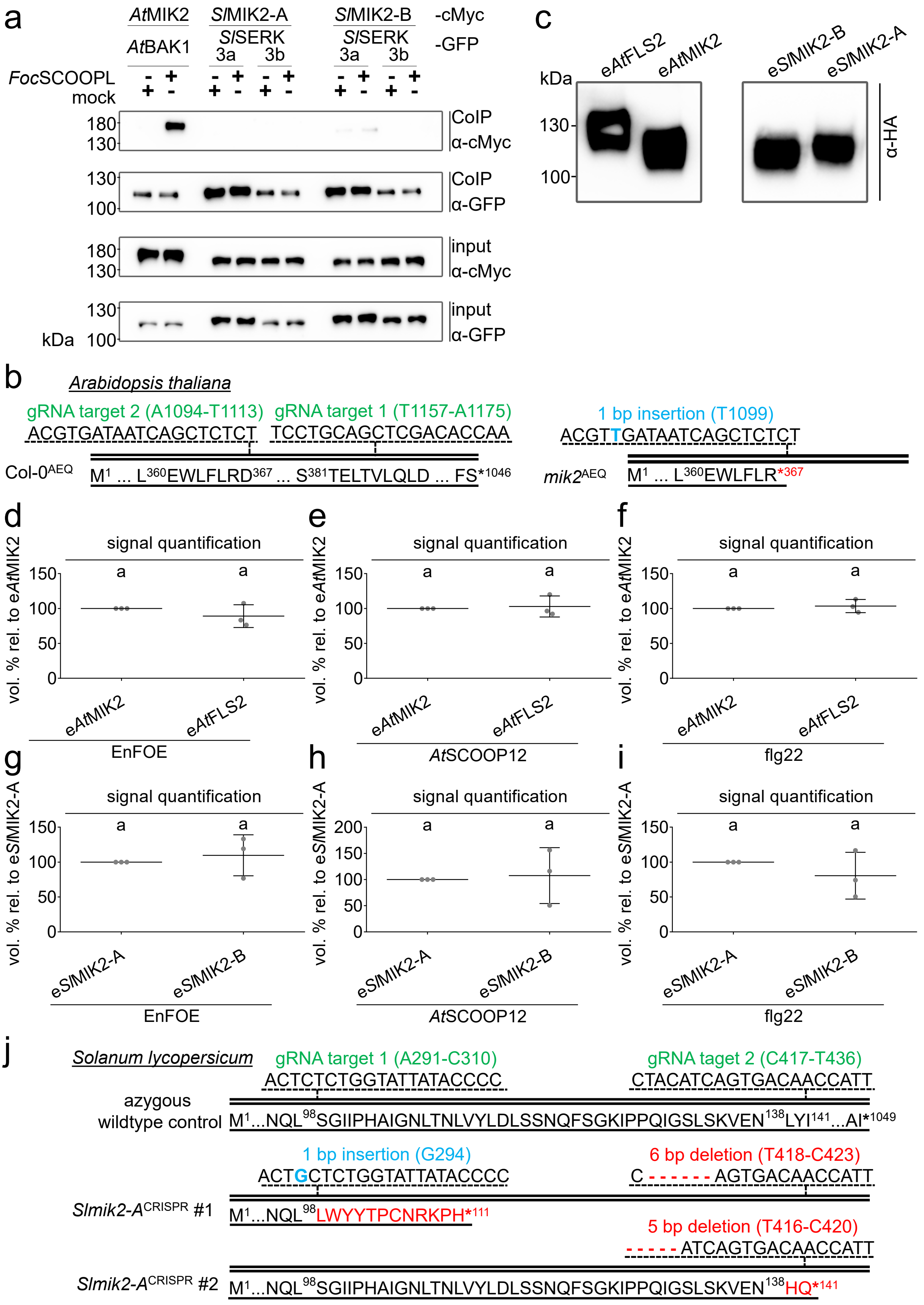


**fig.S4 | Ligand-induced receptor-complex formation, expression of affinity-tagged ectodomains and schematics of *Atmik2*$^{AEQ}$ and *Slmik2-A* CRISPR mutants. (a)** Co-immunoprecipitation of MIK2 and BAK1 proteins from *Arabidopsis thaliana* (*At*MIK2-cMyc, *At*BAK1-GFP) and respective orthologues of *Solanum lycopersicum* (*Sl*MIK2-A-cMyc [splice-variant 1], *Sl*MIK2-B-cMyc, *Sl*SERK3a-GFP, SlSERK3b-GFP) transiently overexpressed in *Nicotiana benthamiana*, 20 min after elicitation with water (mock) or *Foc*SCOOPL$^{FOXB_11846}$ (1 μM). Immunoprecipitation was performed using magnetic anti-GFP beads, western blots were probed with α-cMyc or α-GFP antibodies. Approximate protein sizes are labelled on the left and given in kilodaltons (kDa). **(b)** CRISPR/Cas9-induced mutations in the *Atmik2*$^{AEQ}$ mutant. Amino acid sequence of *At*MIK2 (1045 amino acids) in the Col-0$^{AEQ}$ wildtype is shown on the upper part, positions and sequences of the gRNA target sites are indicated at the top. Introduced insertion mutation (blue letter) in the *mik2*$^{AEQ}$ mutant and the effect on the translated protein sequence are shown on the lower part. The red asterisk indicates the introduced premature stop-codon. Genomic sequences are indicated by the double line, single lines stand for the amino acid sequences, superscript numbers indicate amino acid positions. **(c)** Western blot visualization of C-terminally HA-tagged proteins in eluted fractions corresponding to data of one experiment from fig. 3d-f. Blots were probed with α-HA antibodies. Approximate protein size is marked on the left edge and given in kilodaltons (kDa). **(d-i)** Imaged signals of blots corresponding to **(d)** data of three experiments from fig.3e; **(e)** data of three experiments from fig.3d; **(f)** data of three experiments from fig.3f; **(g)** data of three experiments from fig.3e; **(h)** data of three experiments from fig.3d; **(i)** data of three experiments from fig.3f. Imaged signals were quantified using the built-in feature of the Fusion SL Imager software. Signal intensity was calculated relative to signal of e*At*MIK2 (d-f) or e*Sl*MIK2-A (g-i) within each experiment (set to 100%). Data are pooled from 3 independent experiments, bars indicate mean and standard deviation. Different letters indicate statistically significant differences ($P < 0.05$), unpaired t-test. **(j)** CRISPR/Cas9-induced mutations in two independent *Slmik2-A*$^{CRISPR}$ mutant lines. Amino acid sequence of *Sl*MIK2-A (1048 amino acids) in the azygous wildtype control is shown on the upper part, positions and sequences of the gRNA target sites are indicated at the top. Introduced insertions (blue letter) and deletions (red hyphen) in the two *Slmik2-A*$^{CRISPR}$ mutants and the effects on the translated protein sequences are shown in the middle (line #1) and on the lower part (line #2). Amino acid substitutions are indicated in red, the red asterisks indicate the introduced premature stop-codons. Genomic sequences are indicated by the double line, single lines stand for the amino acid sequences, superscript numbers show amino acid positions.

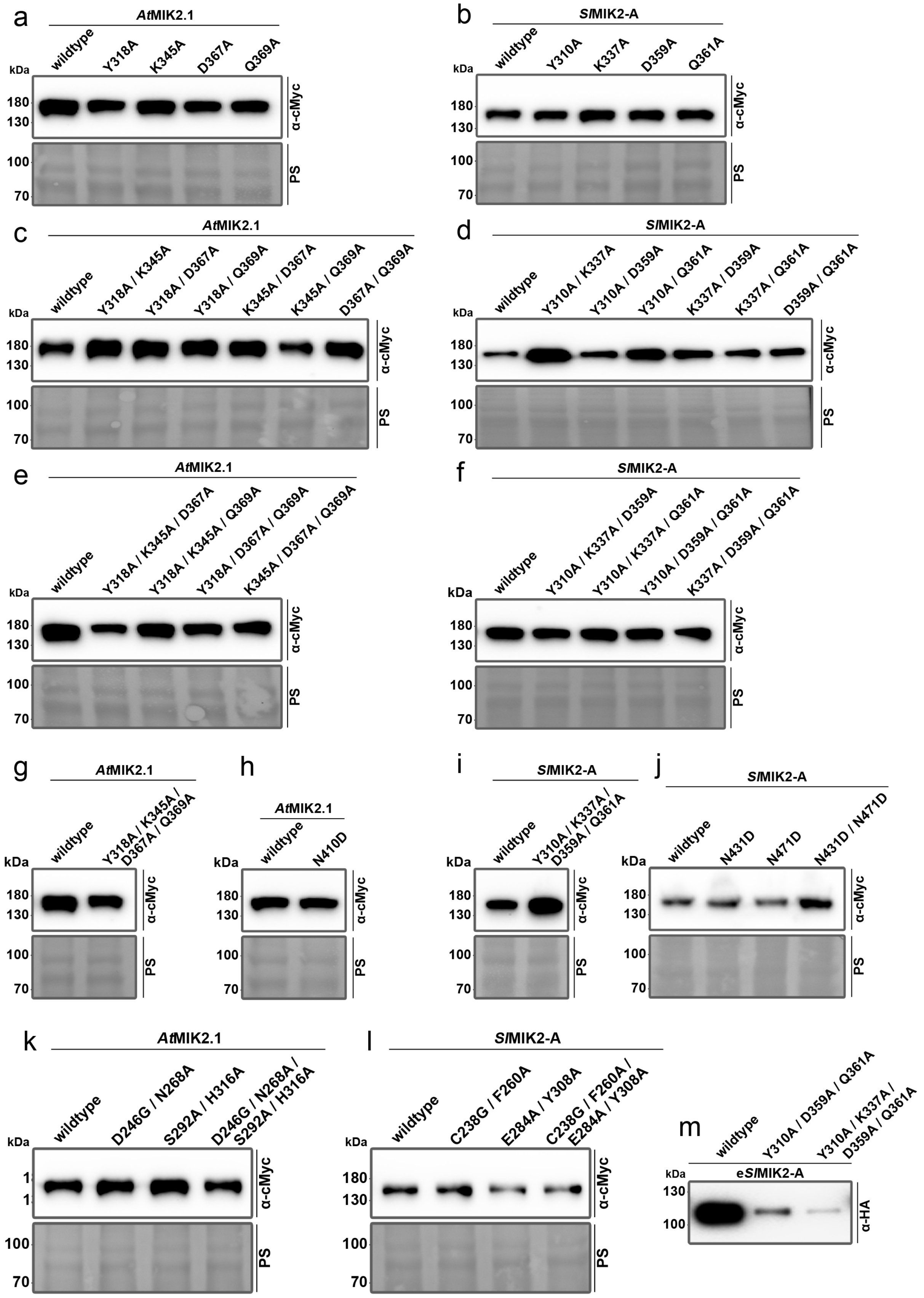


**fig.S5 | Western blot visualization of C-terminally cMyc-tagged proteins.** Blots corresponding to **(a)** data of one experiment from fig.6b, S6b, **(b)** data of one experiment from fig.6d, **(c)** data of one experiment from fig.6c, S6c, **(d)** data of one experiment from fig.6e, **(e)** data of one experiment from fig.S6d-e, **(f)** data of one experiment from fig.S6f, **(g)** data of one experiment from fig.S6g, **(h)** data of one experiment from fig.5e, **(i)** data of one experiment from fig.S6g, **(j)** data of one experiment from fig.5h, **(k)** data of one experiment from fig.5b-c, **(l)** data of one experiment from fig.5d. Blots were probed with α-cMyc antibodies. Approximate protein size is marked on the left edge and given in kilodaltons (kDa). Equal protein loading was confirmed by Ponceau S staining (PS). **(m)** Western blot visualization of C-terminally HA-tagged proteins in eluted fractions from columns lined with wildtype and alanine-substitution mutants of *Sl*MIK-A ectodomains (e*Sl*MIK2-A). Blots were probed with α-HA antibodies. Approximate protein size is marked on the left edge and given in kilodaltons (kDa).

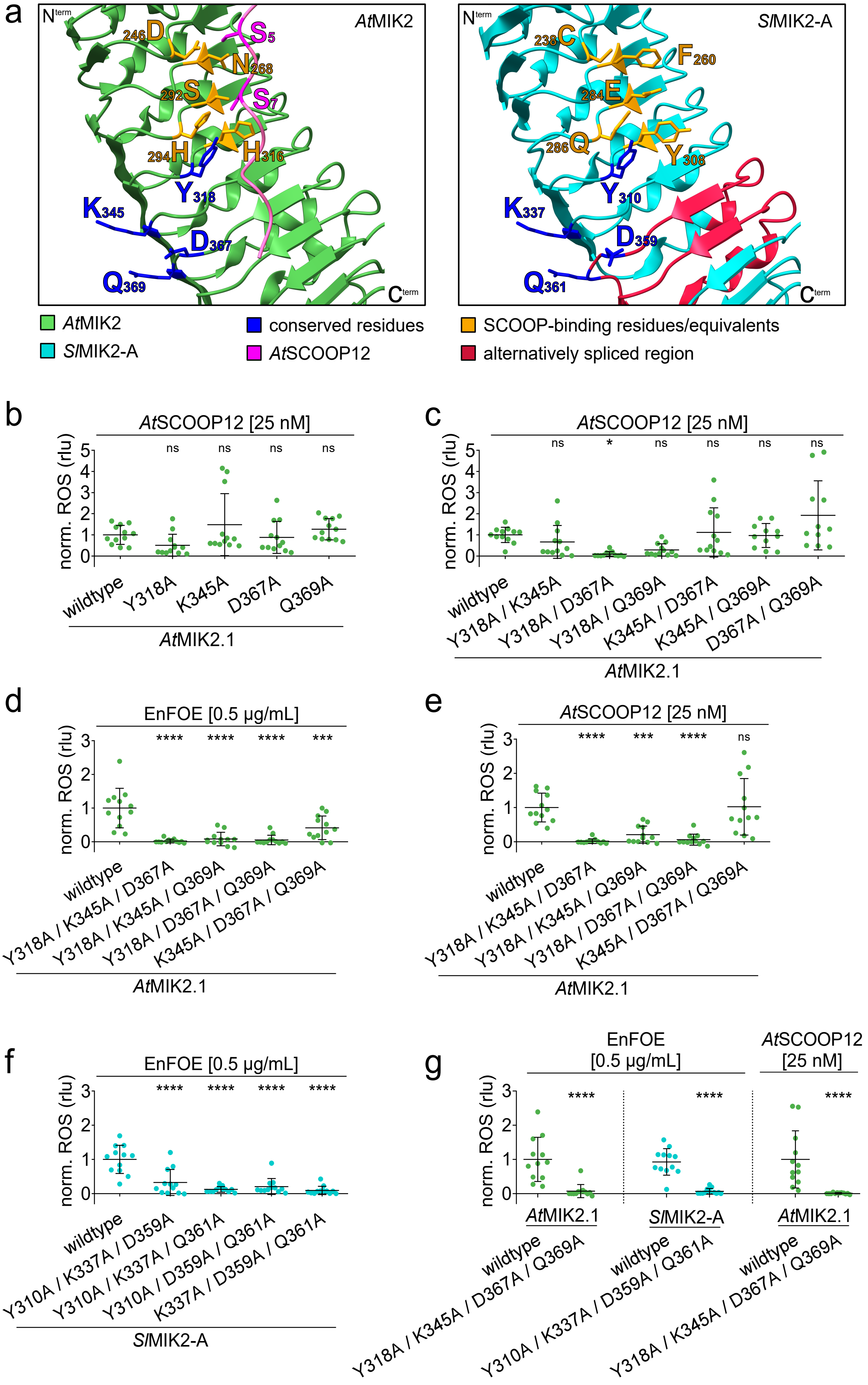
a
AtMIK2
SlMIK2-A
AtMIK2
SlMIK2-A
conserved residues
AtSCOOP12
SCOOP-binding residues/equivalents
alternatively spliced region
b
AtSCOOP12 [25 nM]
norm. ROS (rlu)
wildtype
Y318A
K345A
D367A
Q369A
AtMIK2.1
c
AtSCOOP12 [25 nM]
wildtype
Y318A / K345A
Y318A / D367A
Y318A / Q369A
K345A / D367A
K345A / Q369A
D367A / Q369A
AtMIK2.1
d
EnFOE [0.5 µg/mL]
Y318A / K345A / D367A
Y318A / K345A / Q369A
Y318A / D367A / Q369A
K345A / D367A / Q369A
AtMIK2.1
e
AtSCOOP12 [25 nM]
AtMIK2.1
f
EnFOE [0.5 µg/mL]
Y310A / K337A / D359A
Y310A / K337A / Q361A
Y310A / D359A / Q361A
K337A / D359A / Q361A
SlMIK2-A
g
EnFOE [0.5 µg/mL]
AtSCOOP12 [25 nM]
AtMIK2.1
SlMIK2-A
AtMIK2.1
Y318A / K345A / D367A / Q369A
Y310A / K337A / D359A / Q361A

**fig.S6 | Influence of conserved residues on EnFOE-sensitivity conferred by *At*MIK2 and *Sl*MIK2-A. (a)** Close-up of AlphaFold (AF) structures of *Arabidopsis thaliana At*MIK2 (left, green) and *Solanum lycopersicum Sl*MIK2-A (right, turquoise) ectodomains. *At*MIK2 is shown in complex with *At*SCOOP12 (pink), conserved serine residues (S5 and S7) engaging in the interaction are highlighted. *At*MIK2 SCOOP-binding residues and structural equivalents (same amino acid positions according to structural alignment) on *Sl*MIK2-A are highlighted in orange, residues that are conserved between *At*MIK2 and *Sl*MIK2-A are highlighted in dark blue and respective amino acids and positions are indicated. The region that is alternatively spliced resulting in non-functional *Sl*MIK2-A splice-variant #2 (fig. 2e-g) is marked in red. Orientation of N-termini ($N^{term}$) and C-termini ($C^{term}$) are shown in the corners. **(b-g)** EnFOE- or *At*SCOOP12-induced accumulation of reactive oxygen species (ROS) measured in leaf discs from *Nicotiana benthamiana* transiently overexpressing *Arabidopsis thaliana* MIK2 (*At*MIK2.1) or *Solanum lycopersicum* MIK2-A (*Sl*MIK2-A). Proteins were expressed as wildtype or with indicated amino acid substitutions. Recorded values were normalized to background and mock-treated samples and summed over a period of 60 min (cumulative [cum.] ROS) and are given in relative light units (rlu). To enhance comparability, all individual values were standardized to average values of wildtype controls within each independent experiment (collective values of wildtype average at 1.0). Data are pooled from 3 independent experiments, each with n = 4, bars indicate mean and standard deviation. Asterisks indicate statistically significant differences compared to respective wildtype with **** = $P < 0.0001$, *** = $P < 0.001$, * = $P < 0.05$, ns = not significant with $P > 0.05$, ordinary one-way ANOVA with Dunnett's multiple comparisons test for b-f or unpaired t-test for g.

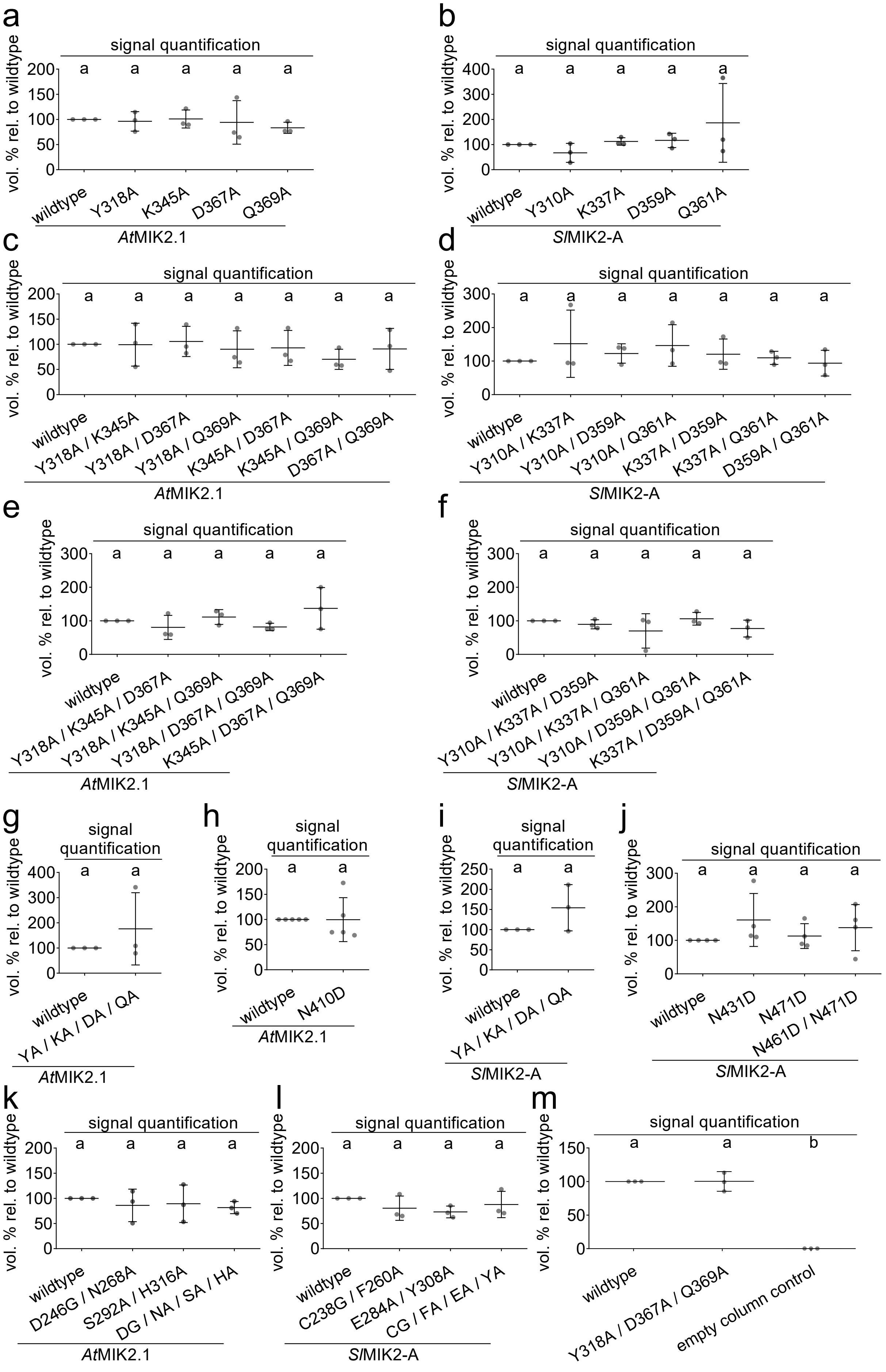
a
signal quantification
vol. % rel. to wildtype
wildtype
Y318A
K345A
D367A
Q369A
AtMIK2.1
b
signal quantification
vol. % rel. to wildtype
wildtype
Y310A
K337A
D359A
Q361A
SlMIK2-A
c
signal quantification
vol. % rel. to wildtype
wildtype
Y318A / K345A
Y318A / D367A
Y318A / Q369A
K345A / D367A
K345A / Q369A
D367A / Q369A
AtMIK2.1
d
signal quantification
vol. % rel. to wildtype
wildtype
Y310A / K337A
Y310A / D359A
Y310A / Q361A
K337A / D359A
K337A / Q361A
D359A / Q361A
SlMIK2-A
e
signal quantification
vol. % rel. to wildtype
wildtype
Y318A / K345A / D367A
Y318A / K345A / Q369A
Y318A / D367A / Q369A
K345A / D367A / Q369A
AtMIK2.1
f
signal quantification
vol. % rel. to wildtype
wildtype
Y310A / K337A / D359A
Y310A / K337A / Q361A
Y310A / D359A / Q361A
K337A / D359A / Q361A
SlMIK2-A
g
signal quantification
vol. % rel. to wildtype
wildtype
YA / KA / DA / QA
AtMIK2.1
h
signal quantification
vol. % rel. to wildtype
wildtype
N410D
AtMIK2.1
i
signal quantification
vol. % rel. to wildtype
wildtype
YA / KA / DA / QA
SlMIK2-A
j
signal quantification
vol. % rel. to wildtype
wildtype
N431D
N471D
N461D / N471D
SlMIK2-A
k
signal quantification
vol. % rel. to wildtype
wildtype
D246G / N268A
S292A / H316A
DG / NA / SA / HA
AtMIK2.1
l
signal quantification
vol. % rel. to wildtype
wildtype
C238G / F260A
E284A / Y308A
CG / FA / EA / YA
SlMIK2-A
m
signal quantification
vol. % rel. to wildtype
wildtype
Y318A / D367A / Q369A
empty column control

**fig.S7 | Quantification of Western blot signals of C-terminally cMyc-tagged proteins.** Imaged signals of blots corresponding to **(a)** data of three experiments from fig.6b; S6b, **(b)** data of three experiments from fig.6d, **(c)** data of three experiments from fig.6c; S6c, **(d)** data of three experiments from fig.6e, **(e)** data of three experiments from fig.S6d-e, **(f)** data of three experiments from fig.S6f, **(g)** data of three experiments from fig.S6g, **(h)** data of five experiments from fig.5e, **(i)** data of three experiments from fig.S6g, **(j)** data of four experiments from fig.5h, **(k)** data of three experiments from fig.5b-c, **(l)** data of three experiments from fig.5d, **(m)** data of three experiments from fig.6f. Imaged signals were quantified using the built-in feature of the Fusion SL Imager software. Signal intensity was calculated relative to respective wildtype controls within each experiment (wildtype signals set to 100%). Data are pooled from 3-5 independent experiments, bars indicate mean and standard deviation. Different letters indicate statistically significant differences (one-way ANOVA, Tukey's multiple comparisons test [or unpaired t-test for g-i]. $P < 0.05$).
